# A lysophospholipase plays role in generation of neutral-lipids required for hemozoin formation in malaria parasite

**DOI:** 10.1101/808709

**Authors:** Mohd Asad, Yoshiki Yamaryo-Botté, Mohammad E. Hossain, Vandana Thakur, Shaifali Jain, Gaurav Datta, Cyrille Y. Botté, Asif Mohmmed

**Author notes:** These authors equally contributed as senior authors (Cyrille Y. Botté, Asif Mohmmed).

## Abstract

Phospholipid metabolism is crucial for membrane dynamics in malaria parasites during entire cycle in the host cell. *Plasmodium falciparum* harbours several members of phospholipase family, which play key role in phospholipid metabolism. Here we have functionally characterized a parasite lysophospholipase (*Pf*LPL1) with a view to understand its role in lipid homeostasis. We used a regulated fluorescence affinity tagging, which allowed endogenous localization and transient knock-down of the protein. *Pf*fLPL1localizes to dynamic vesicular structures that traffic from parasite periphery, through the cytosol to get associated as a multi-vesicular neutral lipid rich body next to the food-vacuole during blood stages. Down-regulation of the *Pf*LPL1 disrupted parasite lipid-homeostasis leading to significant reduction of neutral lipids in lipid-bodies. This hindered conversion of heme to hemozoin, leading to food-vacuole abnormalities, which in turn disrupted parasite development cycle and significantly inhibited parasite growth. Detailed lipidomic analyses of inducible knock-down parasites confirmed role of *Pf*LPL1 in generation of neutral lipid through recycling of phospholipids. Our study thus suggests a specific role of *Pf*LPL1 to generate neutral lipids in the parasite, which are essential for parasite survival.

**Importance:** Present study was undertaken with a view to understand the functional role of a unique lipase (lysophopholipase, *Pf*LPL1) of the human malaria parasite. We utilized genetic approaches for GFP tagging as well as to knock-down the target protein in the parasite. Our studies showed that *Pf*LPL1 associates closely with the lysosome like organelle in the parasite, the food-vacuole. During the blood stage parasite cycle, the food-vacuole is involved in degradation of host haemoglobin and conversion of heme to hemozoin. Genetic knock-down approaches and detailed lipidomic studies confirmed that *Pf*LPL1 protein plays key role in generation of neutral lipid stores in the parasite; neutral lipids are essentially required for hemozoin formation in the parasite, a vital function of the food-vacuole. Overall, this study identified specific role of *Pf*LPL1 in the parasite which is essential for parasite survival.

## Introduction

Malaria remains a major parasitic disease in the tropical and sub-tropical countries that results in∼500,000 to 1 million deaths globally every year(1, 2). There is no current efficient vaccine and the rapid spread of drug resistant parasites strains, even to the front line treatment using artemisinin combinations(3–5), which plead for the identification of new drug-targets and development of new drugs against the disease. Understanding of key metabolic pathways in the parasite that sustain parasite survival within its human host will be critical to identify unique and specific targets in the parasites. The parasite development and division during asexual life cycle in host erythrocytes include dramatic modification of host cell membranous structures (i.e. tubulovesicular network, Maurer’s cleft and knobs) and massive increase of lipid synthesis and membrane biogenesis needed for parasite division and propagation (6, 7). Indeed, the parasites require large amount of lipids to generate new membrane-bound compartments for the assembly of future daughter cells(8–10), for cell signalling event, as well as to generate storage lipids, triacylglycerol (TAG) and cholesteryl-esters (6). Neutral lipids (i.e. TAG, DAG, cholesterol and cholesteryl esters) are suggested to be closely associated with heme detoxification during asexual intra-erythrocytic development by allowing/catalyzing its polymerization into hemozoin (11, 12). Establishment of *P. falciparum* intra-erythrocytic infection is associated with a large increase in the neutral lipid and lipid-associated fatty acid (FA) content in infected red blood cells (iRBCs), suggesting an active lipid metabolism pathway in *P. falciparum* (13). Furthermore, the phospholipids in an infected red blood cell increase up to 500-700% compared to an uninfected erythrocyte. The phospholipid synthesis machinery is present and highly active in the parasite (6), and is initiated by the synthesis of fatty acids (FA), the building blocks and hydrophobic moieties of most membrane and storage lipids. *P. falciparum* possesses a relict non-photosynthetic plastid, i.e. the apicoplast that harbour pathway for *de novo* biosynthesis of fatty acid (FA). This pathway is not essential for asexual stages of the parasite in regular culture conditions(9, 14), but it can be re-activated in adverse lipid composition (7).Furthermore, the parasite is able to scavenge a wide range of FA from the host and specifically relies on the import of C16:0 and C18:1 to divide during blood stages (7, 15). Importantly, the parasite possesses an active *de novo* phospholipid synthesis machinery (Kennedy pathway, CDP-DAG pathway), capable of assembling and synthesizing all lipid precursors (PA, DAG, CDP-DAG) and major phospholipid classes (PC, PE, PS, PI, PG, CL) from precursors (polar heads, Lyso-lipids) scavenged from the host and its environment (16).*P. falciparum* blood stages notably relies on the scavenging of lysophosphatidylcholine (LPC), especially its polar head phosphocholine to fuel the *de novo* synthesis of the major membrane lipid of the parasite, phosphatidylcholine (PC), via the Kennedy pathway and thus maintain asexual division (16, 17). Since the Kennedy pathway utilizes the phosphocholine head group from scavenged LPC to synthesize PC, this phosphor choline head group needs to be released from LPC via an undetermined set of enzymes. Taken together, these data indicate that during blood stages, *P. falciparum* relies on the scavenging of phospholipids/FA/lysolipids from host but the molecular machinery that allow to separate and reassemble the phospholipid building blocks (FA, polar heads, lysolipids) remain to be elucidated. Such catabolism of lipid metabolites from the host must involve phospholipases and lysophospholipases to manipulate the scavenged lipids and generate the proper lipid moieties. Phospholipase-like activities are reported from asexual stage parasites and a phospholipase was shown to play essential role in survival of erythrocytic or liver stages of the parasite (18, 19). *P. falciparum* possess a large family of lysophospholipases (LPLs), which most are remarkably encoded in the sub-telomeric regions of chromosomes. Importantly, biochemical activity of LPL was reported during *P. falciparum* blood stages (20). The LPLs catalyse the hydrolysis of acyl chains from lysophospholipids, the intermediated in metabolism of membrane phospholipids, and therefore LPLs play key role recycling of lipids.

In the present study, we have carried out detailed functional characterization of one of the *P. falciparum* LPL, which we named as *Pf*LPL1 (also see supplementary Table S1). Owing to the technical challenges for the disruption of essential genes in *P. falciparum* blood stages and obtain the respective stable transgenic parasite lines, novel tools have been developed for gene functional analysis, including the conditional disruption-degradation domain-based regulated fluorescence tag (21). We used this system for both endogenous localization and transient down-regulation of PfLPL1during the parasite blood stages. The GFP targeting approach demonstrated that the enzyme is associated with vesicles, which traffic from the parasite periphery (i.e. plasma membrane) towards a compartment accumulating neutral lipids in the vicinity of the food vacuole. To confirm its functional role, we performed lipidomic analyses, which showed that parasites lacking *Pf*LPL1displayed fluctuations in phospholipid levels, significant reduction of triacylglycerol (TAG), and increased levels of DAGs. Analysis of the content and formation of hemozoin showed that parasite lacking *Pf*LPL1 were deficient in the normal generation of hemozoin. Taken together, our data suggest that *Pf*LPL1 is involved in phospholipid catabolism to generate precursors required for neutral lipids (TAG) synthesis for heme-detoxification, which are essential for parasite survival.

**Figure 1:**
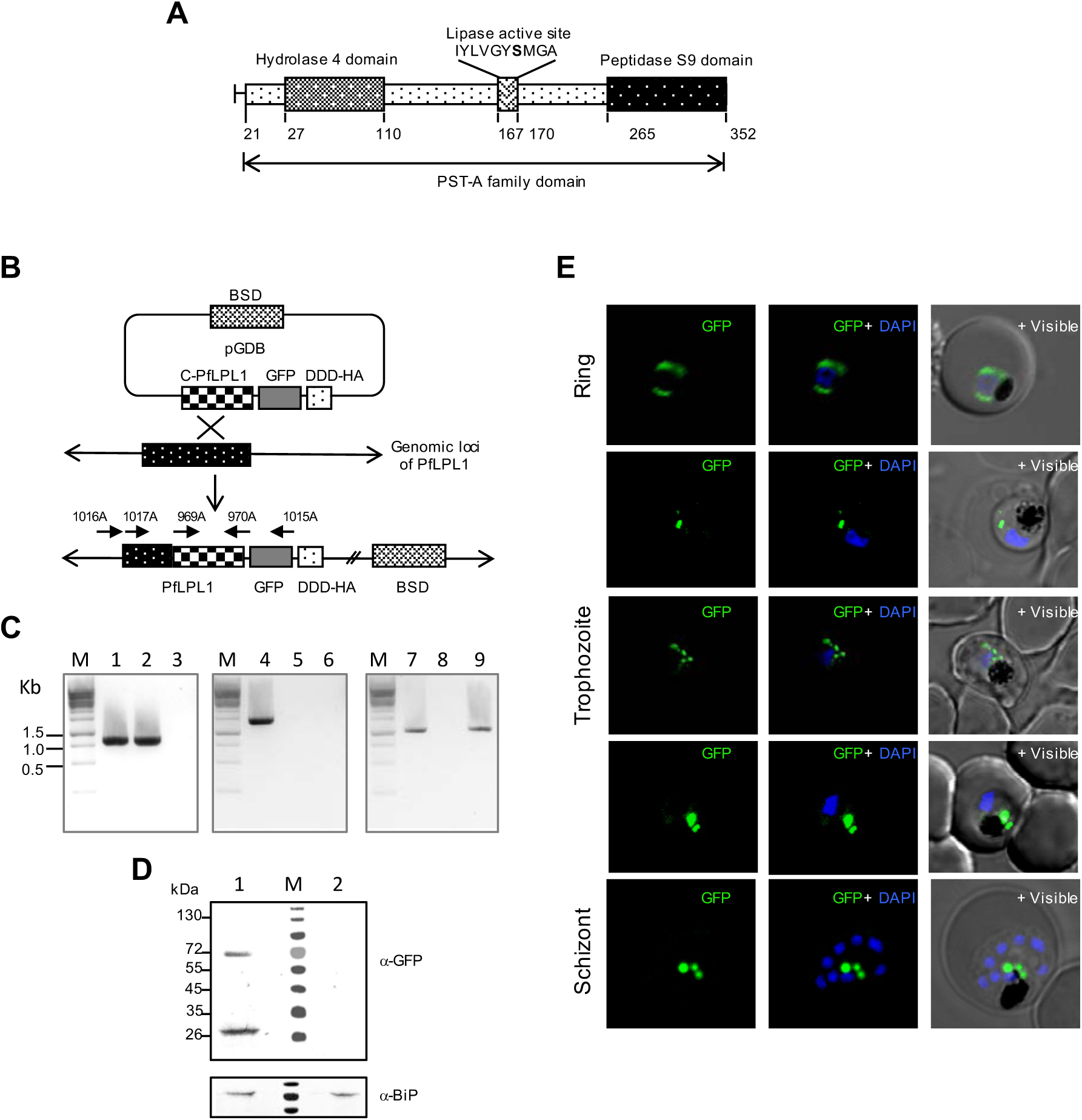
Integration at RFA-tag (DDD-GFP) with *pfLPL1* gene locus and localisation of *Pf*LPL1-RFA fusion protein in transgenic parasites. **(A)** Schematic representation of *Pf*LPL1showing the hydrolase domain and putative peptidase domain. Locations of conserved residues of putative lipase/serine-hydrolase active site are also marked. This fixed pattern GXSXG including the active serine residue is conserved throughout all putative serine hydrolases. **(B)** Schematic diagram showing the strategy used to incorporate the regulatable fluorescent affinity (RFA) tag at the 3’ end of endogenous locus of *pfLPL1* through single cross over homologous recombination. The pGDB vector contains a blasticidin resistance cassette (BSD) for positive selection. The C-terminal fragment of *pfLPL1* gene was cloned in frame with RFA-tag which consists of *E. coli* dihydrofolate reductase degradation domain (DDD) with GFP and HA sequences. Schematic diagrams of the *pfLPL1*genomic loci before and after the homologous recombination are shown. **(C)** PCR based analysis of transgenic parasite cultures after drug cycling to show integration of RFA-tag in the endogenous loci. Primers combinations: 842A-354A (lane 1); 838A-839A (lane 2); and 841A-842 (lane 3). Primers locations in the gene loci or vector construct are indicated in the schematic diagram. **(D)** Immunoblot analysis using anti-GFP antibodies and trophozoite stage transgenic parasites expressing PfLPL1-RFA. A band of ∼70 kDa, representing the fusion protein, and another band of ∼26kDa, representing GFP, are detected in the transgenic parasites (lane 2), but not in the wild-type parasites (lane 1). Lower panel shows a parallel blot probed with anti-BiP antibodies to show equal loading. **(E)** Localization of *Pf*LPL1-RFA fusion protein in transgenic *P. falciparum*parasites. Fluorescent microscopic images of live transgenic parasites at ring, trophozoite, and schizont stages. The parasite nuclei were stained with DAPI and slides were visualized by confocal laser scanning microscope. In ring stages, the GFP-fluorescence was observed around the nucleus; in early trophozoite stage small vesicles are present near the parasitophorous vacuole and in the parasite cytosol. In mature stages, late trophozoite stage and schizonts, GFP-fluorescence was observed in large vesicular structure in close association with the food-vacuole.

## Results

### Endogenous tagging of *pfLPL1* gene and localization of *Pf*LPL1-GFP fusion protein in transgenic parasites

To study the expression, localization and essentiality of the PfLPL1in the parasite, we used the regulated fluorescent affinity (RFA) tag system for C-terminal tagging of the native gene (21). The RFA tag includes GFP in frame with the *E. coli* degradation domain (DDD). The endogenous *pfLPL1*gene was tagged with RFA-tag by single-crossover homologous recombination (Figure 1A, B, C), so that the expression of fusion protein was under the control of native promoter. The transgenic parasites showed expression of *Pf*LPL1-RFA fusion protein of ∼70kDa, migrating at the expected predicted size for the endogenously DDD-GFP tagged protein. Such band was not detected in wild-type 3D7 parasites (Figure 1D).

These transgenic parasites were studied for localization of the *Pf*LPL1-RFA fusion protein by confocal microscopy. In early stages of the parasite, rings and young trophozoites, fluorescence of the GFP fusion protein was localized around the nucleus, which corresponds to the ER region (Figure 1E, Figure 2A). Indeed, staining by ER-tracker showed overlap with the GFP fluorescence in these parasites (Figure 2A). In trophozoite stages, the GFP fluorescence was observed in distinct foci/vesicle-like structures in the parasite. In mid-trophozoite stages, these vesicles were observed near the parasite periphery towards parasitophorous vacuole (Figure 1E). In later stages a number of vesicles were observed further in the parasite cytosol and in close proximity of the food vacuole. In the late-trophozoites and schizont stages the GFP fluorescence accumulated near the food vacuole in 1 or 2 large structure (Figure 1E, Figure S3A).

**Figure 2:**
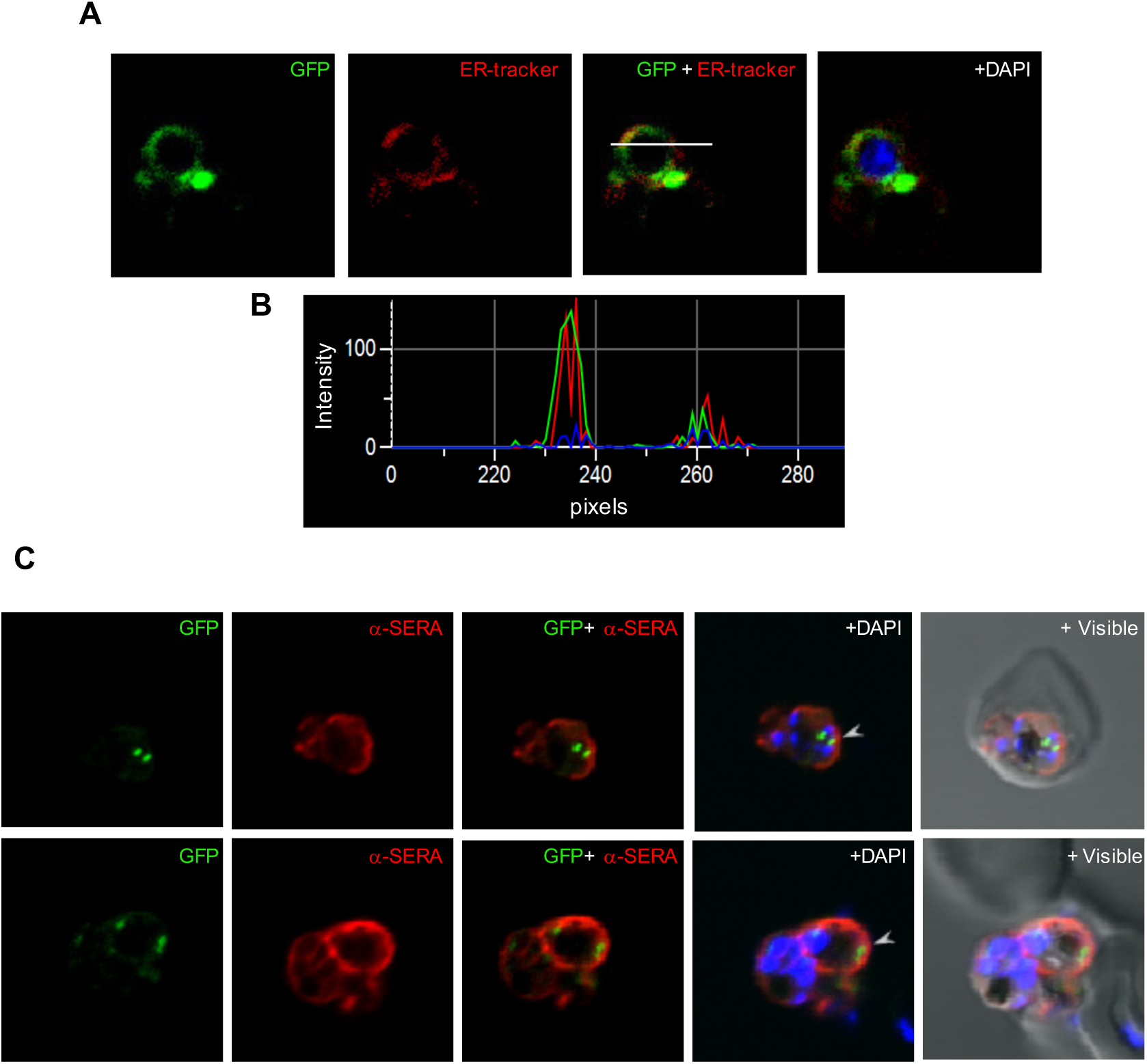
Sub-cellular localization of *Pf*LPL1-RFA fusion protein in transgenic parasite. **(A)** Fluorescent microscopic images of live transgenic parasites at ring stages stained with ER-Tracker (red), the parasite nuclei were stained with DAPI (blue) and slides were visualized by confocal laser scanning microscope. **(B)** A histogram plot for the fluorescence intensity analysis for GFP (green) and ER-tracker (red); the image was analysed by NIS elements software across the stained parasite (marked with white line). **(C)**Fluorescence images of transgenic parasites at trophozoite stages immuno-stained with anti-SERA antibody (red). The PfLPL1-GFP labeled vesicles are seen close to the parasite boundaries inside the parasite cytosol (marked with arrowhead). Parasite nuclei were stained with DAPI (blue) and the parasite were visualised by confocal laser scanning microscope.

### *Pf*LPL1 containing vesicles traffic from parasite periphery towards vicinity of food-vacuole to develop into a multi-vesicular like structures in asexual parasite stages

To further investigate these intriguing and dynamic localization patterns, a series of cellular staining and immuno-staining were carried out and analyzed by confocal microscopy. The transgenic parasites were immuno-stained with anti-SERA, as a marker of the parasitophorous vacuole (22). The anti-SERA antibody clearly stained around parasite periphery defining the parasitophorous vacuole. The *Pf*LPL1-GFP labeled vesicles were seen in close proximity with the anti-SERA staining but did not overlap with it, suggesting that the GFP-vesicles are present at parasite boundary within the parasite and turned towards the parasite cytosol (Figure 2B). Further, we labeled the parasite membranes by a non-specific lipid membrane probe BODIPY-TR ceramide, which can labelparasite plasma membrane, parasitophorous-vacuole membrane, tubulo-vesicular membranes and food vacuolemembrane. The *Pf*LPL1-GFP labelled vesicles were observed in association with parasite boundarybut clearly distinct from membrane labelling(Figure 3A, Figure S4A). A three-dimensional reconstruction based upon Z-stack images of these parasites confirmed the close proximity (without direct association) of these vesicles with the parasite membrane (Figure 3B). Importantly, initial observations suggested that these *Pf*LPL1-GFP vesicles were originating from the parasite periphery; traffic in parasite cytosoland then accumulating in close vicinity to the food vacuole (Figure 3A, B and S4).

**Figure 3:**
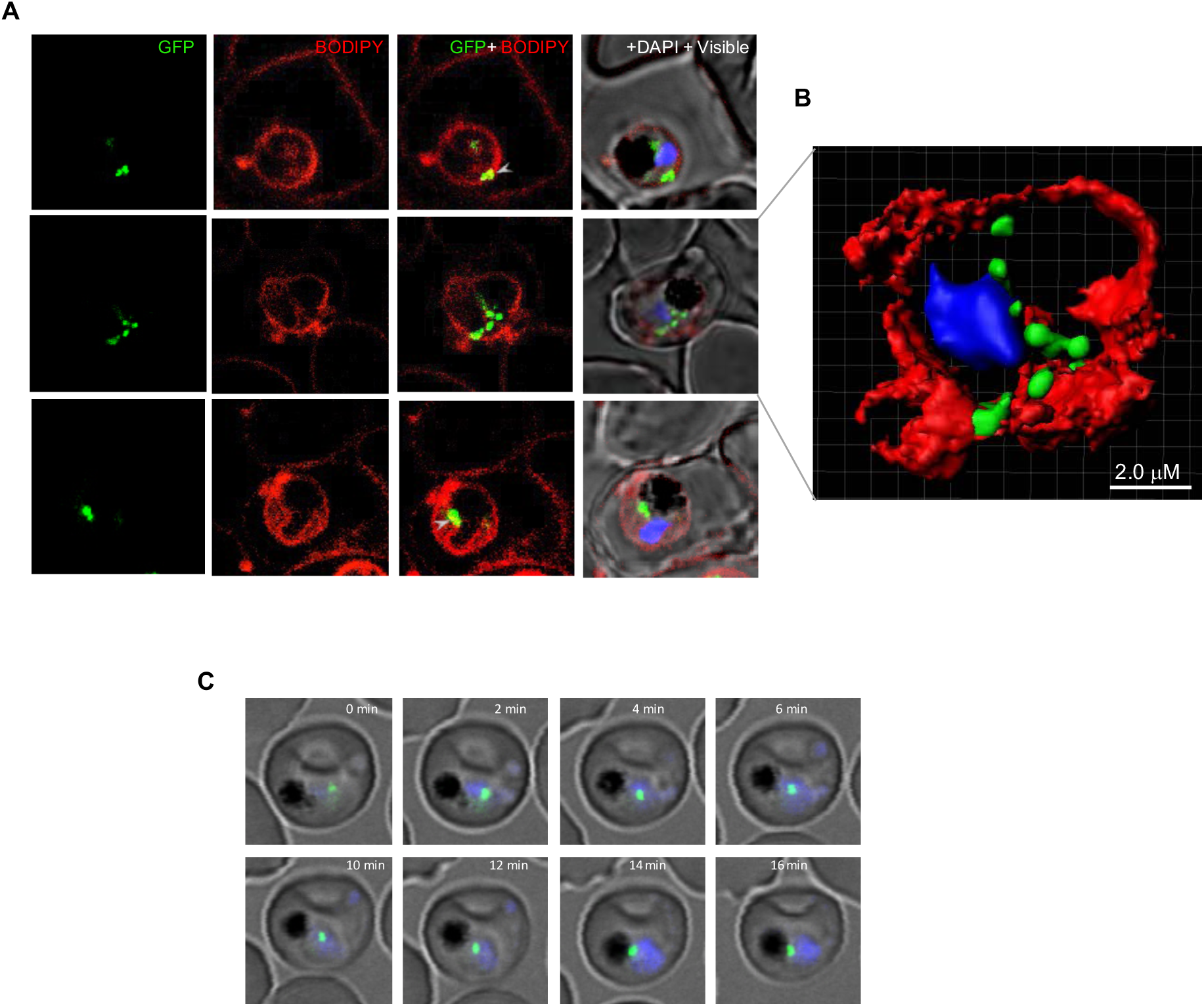
*Pf*LPL1 resides in vesicles which associate parasite membrane and with food-vacuole. **(A)** Fluorescence images showing labelling of membranes in *P. falciparum*infected RBCs and localization of PfLPL1. Trophozoite stage transgenic parasites expressing *Pf*LPL1-RFA were stained with BODIPY-TR ceramide (red), the parasite nuclei were stained with DAPI (blue) and visualized by confocal laser scanning microscope. Small GFP foci of the *Pf*LPL1-RFA fusion protein were observed near parasite boundary (panel 1, marked with arrowhead), these foci showed closed association with fluorescence by BODIPY-TR labelled parasite membrane. In some parasites, the GFP vesicles are seen in parasite cytosol (panel 2) and in close association with food-vacuole (panel 2 and 3). **(B)** A three-dimensional reconstruction of series of Z-stack images (corresponding to panel 2 in A) using IMARIS software. Small GFP vesicles are present juxtaposed to the parasite-membrane, in parasite cytosol and close to the food-vacuole. **(C)** Time-lapse microscopy of *Pf*LPL1-RFA expressing transgenic parasites showing localization and migration GFP labeled vesicles. Sequential images of a trophozoite stag parasite over a time interval of 16 min showing trafficking of a GFP labeled vesicle fin the parasite cytosol which subsequently gets associated at the boundary of food-vacuole (having dark hemozoin).

To further understand the trafficking of PfLPL1in the parasites, we determined the localization and movement of *Pf*LPL1-GFP tagged vesicles in the transgenic parasites by time-lapse confocal microscopy. Observations of trophozoite stage parasites expressing PfLPL1-GFP using time-lapse imaging showed different phases of vesicles development and movement in the parasites (Figure 3C). In this set of live-imaging, *Pf*LPL1-GFP labeled vesicular structures first appears near the parasite plasma membrane and throughout the parasite cytosol (0-4min). The vesicles then seem to appear as a single GFP-labelled structure, which gradually migrate towards the food vacuole (6-14min) and subsequently localizes in close proximity to the food-vacuole (Figure 3C). In another set of images, a number of GFP labeled vesicles are seen originating from the parasite membrane. These vesicles subsequently migrated in the parasite cytosol and ultimately culminated as a single structure near the food-vacuole (Figure S3B). Overall these results show migration of *Pf*LPL1-GFP containing vesicles in the parasite cytosol and their culmination into a multi-vesicular like body near food-vacuole.

To further ascertain the association of *Pf*LPL1 with vesicular structures, we performed immuno-electron microscopic studies of the transgenic parasite using anti-GFP antibodies. In early ring stages, immuno-labelling of *Pf*LPL1-GFP was observed in the endo-membrane system of the parasite (nuclear envelope-ER area) (Figure 4A, B). In trophozoite stages, the immuno-labelling was observed in small membrane bound vesicles of ∼100nm size, and some of these labeled vesicles were seen originating near the parasite plasma membrane (Figure 4C), whereas some vesicles were observed distributed in the parasite cytosol (Figure 4C). Mature trophozoite stages displayed large clearly labeled membrane bound structures (∼200-300nm) that were observed closely associated with the food-vacuole (Figure 4D, E). Specificity of the observed structures for *Pf*LPL1-GFP vesicles was confirmed with immuno-EM controls conducted with secondary antibodies alone or pre-immune mice sera.

**Figure 4:**
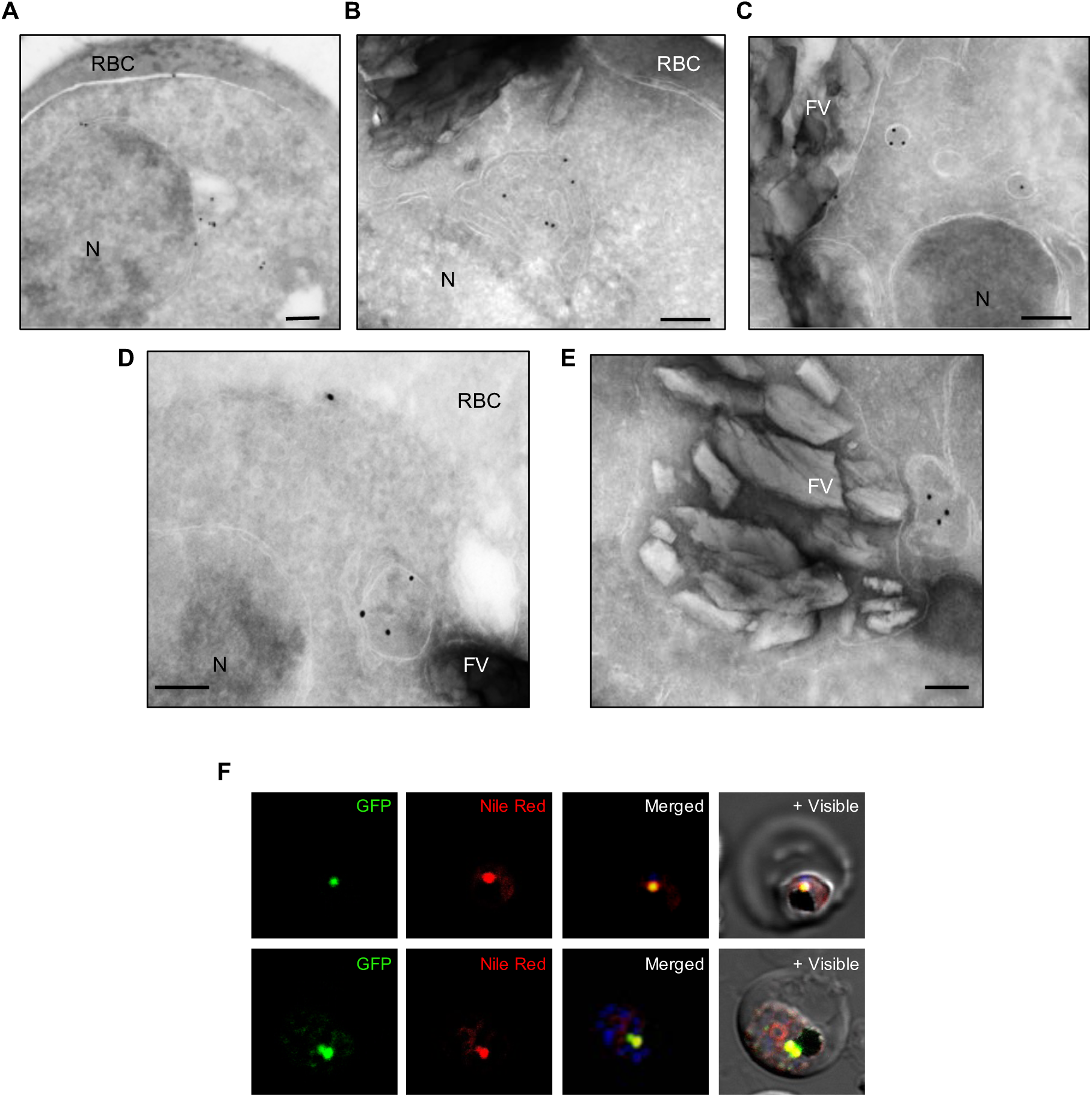
**(A-E)** Localization of *Pf*LPL1 by immune-electron microscopy. Ultra-thin sections of transgenic *P. falciparum* parasites expressing PfLPL1-RFA were labelled with anti-GFP antibody and gold labelled secondary antibody: **(A)** In young parasites, labelling was observed around nucleus in the endoplasmic reticulum; **(B)** in some young parasite labelling was in the putative ER exit site; **(C)** In trophozoite stages, labeling was observed in small vesicles (<100nm) in the parasite cytosol; **(D, E)** in mature parasites, labeling was observed in large vesicular structure (>200nm) localized in close association with the food-vacuole (having characteristic hemozoin). Scale bar = 200nm. **(F)** *Pf*LPL1associate with neutral lipid storage body near food vacuole. Fluorescence images of trophozoites stage transgenic parasites expressing PfLPL1-RFA stained with Nile Red, a neutral lipid staining dye. The large GFP labelled structure close to the food vacuole (having dark hemozoin) showed staining with Nile Red, whereas other GFP vesicles in cytosol were not stained with Nile Red (marked with arrow-head). The parasite nuclei were stained with DAPI (blue) and parasites were visualized by confocal laser scanning microscope.

Overall localization analyses strongly indicate that: (1) *Pf*LPL1 is associated with ER in early ring stages; (2) it gets further trafficked from ER towards parasite periphery during early trophozoite stage and gets associated with cytosomal-like vesicles, which are developing during uptake of content of parasitophorous vacuole/host cytosol; (3) the *Pf*LPL1 protein subsequently traverses along with these cytosomal like vesicles in parasite cytosol; and (4) these vesicles get associated/fuse with each other in the form a large structure near parasite food vacuole in mature parasite stages.

### *Pf*LPL1 associates with neutral lipid storage site (lipid bodies) in the parasite

Since PfLPL1 is expected to generate free fatty acid (FA) from lyso-lipids, we assessed any potential association of the protein with neutral lipid bodies or lipid storage that could accumulate such FA. We determined localization of *Pf*LPL1with respect to lipid storage vesicles using Nile Red. Nile Red is a hydrophobic probe that can stain lipid storages in cells and concentrate in stores of neutral lipids such as TAG (23). The *Pf*LPL1-RFA parasites were stained with Nile Red and visualized with confocal microscopy. In trophozoite stage parasites the Nile Red staining showed a single intensely fluorescent spot closely associated with food vacuole (Figure 4F). Similarly, 1-2 of these intensely labeled spots representing neutral lipid rich structures were observed in late schizont stages. The Nile Red stained lipid bodies showed complete overlap with GFP fluorescence near the food vacuole, suggesting that the large vesicular structures that contain the *Pf*LPL1near food vacuole are the neutral lipid storage bodies near the parasite food vacuole (Figure 4F).

To ascertain that GFP-tag is not influencing the localization of native protein in the parasite, we generated transgenic parasite where *pfLPL1*gene was tagged at the C-terminal to express the native gene fused with HA-DD tag (see Supplementary data; Figure S5). Localization of the fusion protein was assessed by immuno-staining with anti-HA antibodies and Nile red staining. All previous results obtained with *Pf*LPL1-GFP fusion parasite lines were confirmed in the *Pf*LPL1-HA-DD lines: *Pf*LPL1 protein was localized in vesicular structures, these vesicles were also observed near the parasite periphery in early parasite stages, and in later stages a number of vesicles were seen in the parasite cytosol and in close proximity of the food vacuole (Figure S6). In late-trophozoites and schizont stages the protein was found to be accumulating near the food vacuole in 1 or 2 large multi-vesicular like structure (Figure S6A, B). These structures showed clear overlap with Nile Red staining in the parasites (Figure S6C).

Overall, results of localization studies by GFP-tagging/HA-tagging and Nile Red staining show that the *Pf*LPL1 is associated with neutral lipid storages and thus likely to be involved in generation of neutral lipids in the parasite.

### Selective degradation of *Pf*LPL1 inhibits parasite growth and disrupts the intra-erythrocytic developmental cycle

To understand the functional significance of *Pf*LPL1 and its possible involvement in neutral lipid synthesis/storage we utilized the RFA tagged mediated selective degradation of *Pf*LPL1 in the transgenic parasites. Selective degradation of RFA-tagged protein in absence of TMP was assessed by quantifying fluorescent cells in parasite cultures, which were grown in presence or absence of TMP. As shown in Figure 5A, absence of TMP caused significant reduction in fluorescent intensity of the parasites and reduced ∼90% of fluorescent parasites as estimated by flow-cytometry based analysis. Effect of this inducible knock-out of *Pf*LPL1 (*Pf*LPL1-iKO) was assessed on parasite growth *in vitro* by estimating development of new ring stage parasites for 3 cycles. In *Pf*LPL1-iKO set, the parasite growth was significantly reduced (∼60%) as compared to control parasite culture (Figure 5B). Growth inhibition studies using *Pf*LPL1-HA-DD lines showed much enhanced effect; inducible knock-down of *Pf*LPL1 in iKO set showed ∼80% growth inhibition as compared to control set (Figure S7). These results suggest that *Pf*LPL1 plays a critical role for the intra-erythrocytic development of *P. falciparum*.

**Figure 5:**
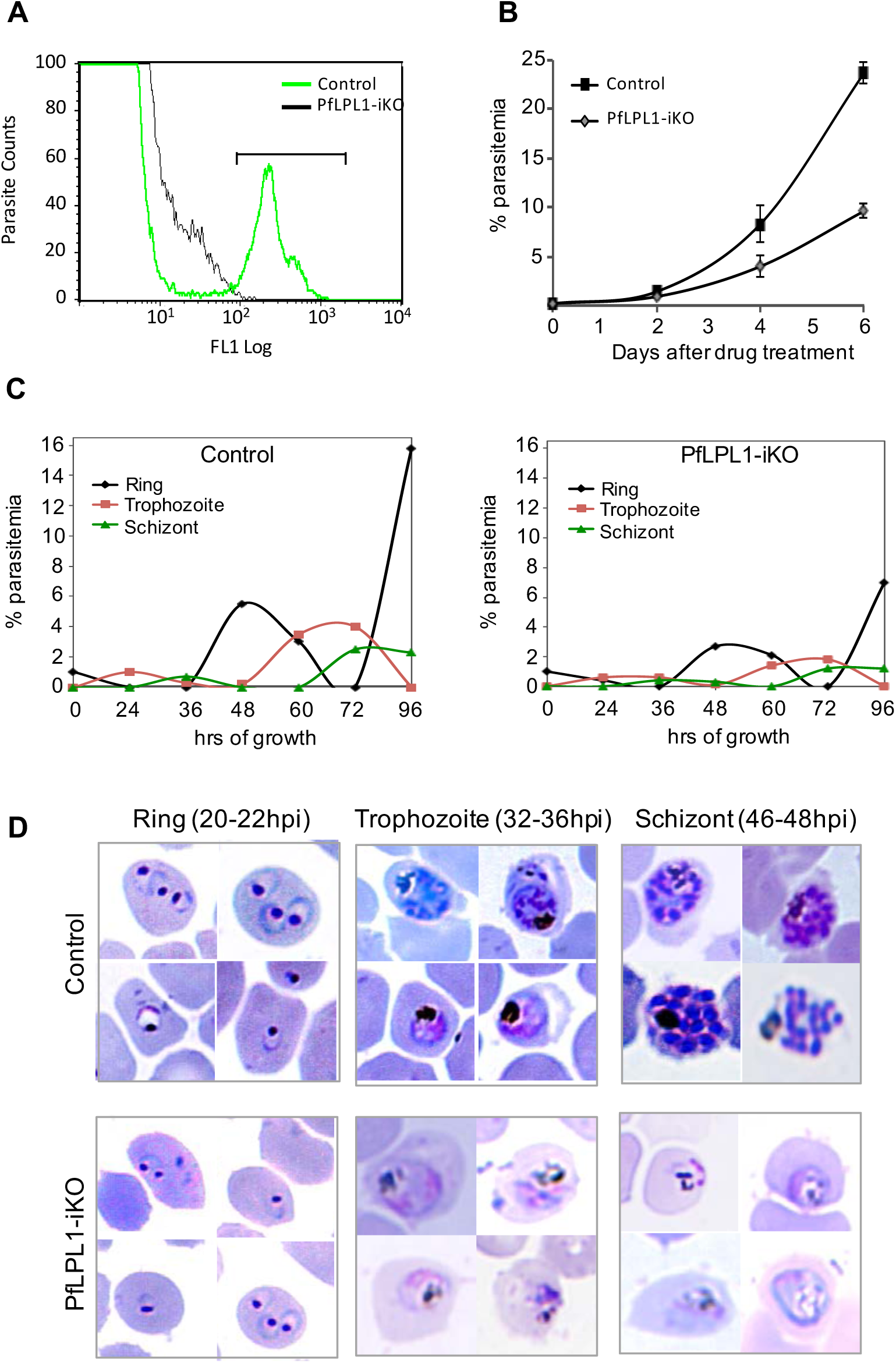
Inducible knock-down of *Pf*LPL1 protein in the transgenic parasites and its effect on growth and development asexual cycle of the parasites. **(A)** Flow cytometry histogram showing reduction in population of GFP-labelled parasites in transgenic parasite cultures after PfLPL1 inducible knock-down (grown in absence of TMP) as compared to control parasites (grown in presence of TMP). **(B)** Tightly synchronized ring stage parasite culture (0.2% parasitemia) of transgenic parasites were grown with or without TMP (Control and *Pf*LPL1 inducible knock-down, *Pf*LPL1-iKO, respectively), and their growth was monitored for three cycles by estimating total parasitemia at 48, 96 and 144h. **(C)** Graphs showing parasite stage composition at different time points (0-120h) in parasite culture from control and *Pf*LPL1-iKO sets. **(D)** Giemsa stained images of parasites showing effect on parasite morphology at different time points (0-48h) in parasite culture control and *Pf*LPL1-iKO sets.

To study the effect of selective degradation of *Pf*LPL1on parasite development and morphology, synchronized ring stage parasites in *Pf*LPL1-iKO and control sets were analyzed for several intra-erythrocytic cycles. In control set, during each intra-erythrocytic cycle the parasites developed from ring to trophozoites to mature schizonts and subsequently merozoites released from these schizonts invaded new erythrocytes, which effectively increased the total parasitemia about 6 times (Figure 5C, D). However, in the *Pf*LPL1-iKO set, development of parasite from trophozoites to schizont was hindered (Figure 5C, D). The *Pf*LPL1-iKOset also showed morphological abnormalities during development. The mature trophozoite stages showed abnormal food-vacuoles without distinct hemozoin crystals as compared to control parasites that showed clear dark hemozoin pigment (Figure 5D). This resulted in reduction in parasitemia in next cell cycle. The loss of parasitemia was detected again in subsequent cycles leading to significant reduction in overall growth of the parasites.

### Knock-down of *Pf*LPL1 resulted in reduction of food-vacuole associated neutral lipids

To understand possible role of *Pf*LPL1 in biogenesis and storage of neutral lipids, we assessed any effect of selective knock-down of *Pf*LPL1 levels in the parasite on accumulation of neutral lipids. Synchronized ring stage *Pf*LPL1-RFA parasites in control and *Pf*LPL1-iKO sets were grown for 24-30h to develop into trophozoite stages, subsequently levels of *Pf*LPL1-GFP and neutral lipids levels were assessed in these parasites. Flow cytometry-based analysis as well as fluorescence microscopy showed significant reduction of GFP fluorescence in the parasites (Figure 6A, C) and reduction in Nile Red staining of the lipid storage bodies (Figure 6B, C) after downregulation of *Pf*LPL1 (*Pf*LPL1-iKO), as compared to control parasites. Inducible knock-down studies using *Pf*LPL1-HA-DD lines showed similar effect on reduction in neutral lipid storage body (Figure S7C).

**Figure 6:**
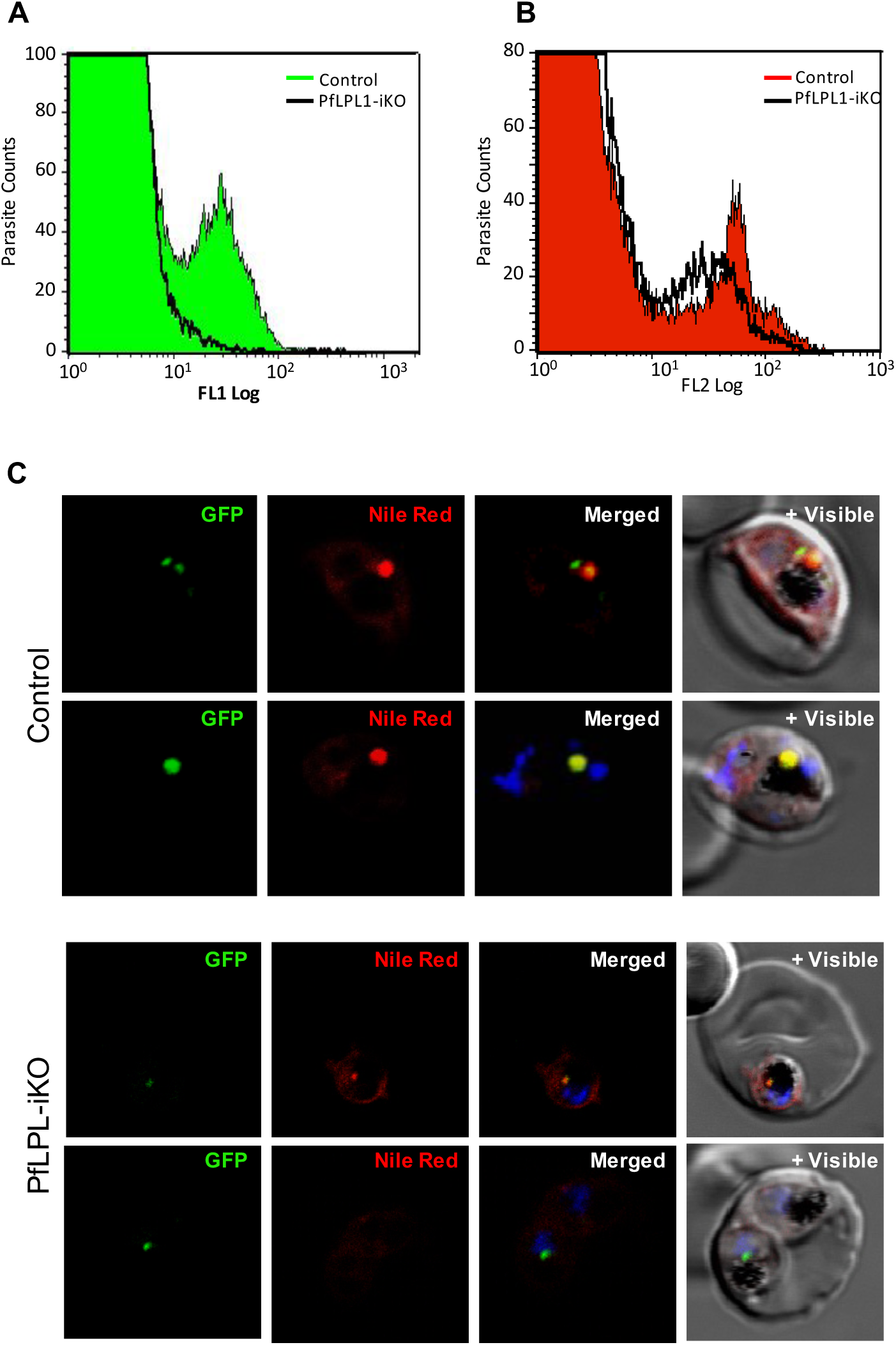
Inducible knock-down of *Pf*LPL1 protein in the transgenic parasites and its effect on the development of neutral lipid body. **(A and B)** Synchronous transgenic parasites at ring stages were grown till late trophozoite stages in control and *Pf*LPL1-iKO sets, stained with Nile red and analysed by flow cytometry. Flow cytometry histogram showing concomitant reduction in GFP fluorescence (FL-1) and Nile red labelling (FL-2) in parasites after *Pf*LPL1-iKO as compared to control parasites. **(C)**Fluorescence images of trophozoites stage transgenic parasites in *Pf*LPL1-iKO set, showing the reduction in Nile red fluorescence intensity and loss of GFP-fluorescence as compared to control parasite. The parasite nuclei were stained with DAPI (blue) and parasites were visualized by confocal laser scanning microscope.

### Downregulation of *Pf*LPL1 results in major changes in the parasite lipid composition, and reveals its potential role for TAG synthesis

To determine the role of *Pf*LPL1 at the lipid synthesis levels and its putative associated function with neutral lipid storage, we conducted mass spectrometry-based lipidomic analyses on lipid extracted from synchronized *Pf*LPL1-RFA ring stage parasites grown further for24h from control and inducible knock-down culture sets. Analysis of FA from total lipid extraction showed no significant changes in the FA composition of parasites from control and *Pf*LPL1-iKO (Figure 7A), where the main FA species are C16:0, C18:0 and C18:1 as previously reported(24, 25). However, analysis of the parasites phospholipid composition revealed significant changes in *Pf*LPL1-iKO. Indeed, relative abundance of phosphatidic acid (PA) and phsophatidylethanolamine (PE) were significantly reduced in the knock-down cultures, whereas relative abundance of both phosphatidylcholine (PC) and phosphatidylserine (PS) were significantly increased (Figure 7B). Furthermore, analysis of the content of major neutral lipids making the bulk of lipid bodies composition, i.e. diacylglycerol (DAG) and TAG, showed that the relative abundance of DAG was significantly increased whereas the one from TAG was significantly reduced after *Pf*LPL1 knock-down (Figure 7C). Together these results correlate with our phenotypic analysis that *Pf*LPL1 is involved in the generation of neutral lipids, and more particularly TAGs, but also impacts the content of the parasite phospholipid.

**Figure 7:**
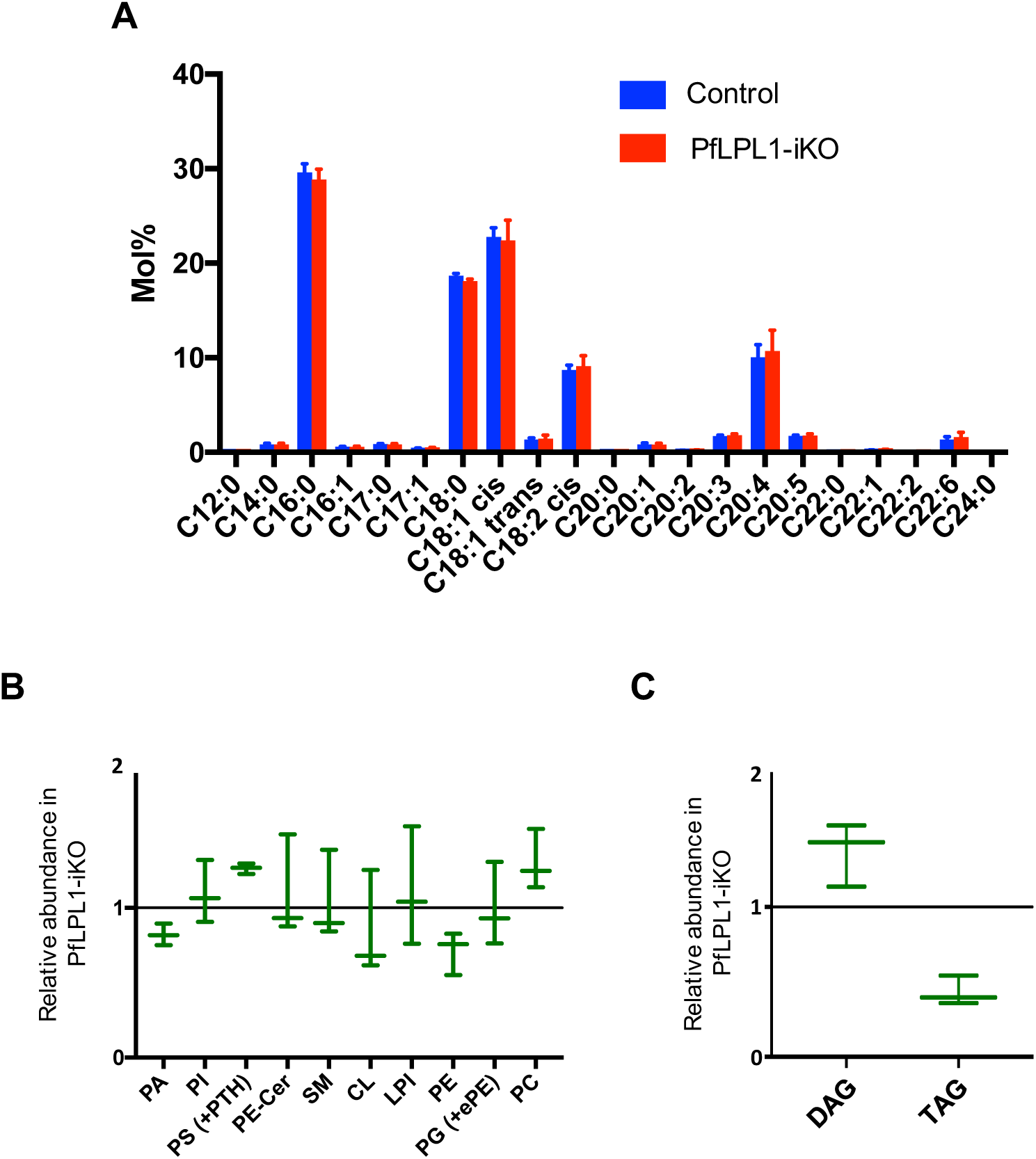
Inducible knock-down of *Pf*LPL1 protein in the transgenic parasites and its effect on lipid homeostasis. Synchronous transgenic parasites at ring stages were grown till late trophozoite stages and lipids compositions were assessed by mass spectrometry-based lipidomic analyses from *Pf*LPL1-iKO and control sets. **(A)** Analysis of the parasites phospholipid composition showing downregulation of phosphatidic acid (PA) and phsophatidylethanolamine (PE) and upregulation of phosphatidylcholine (PC) and phosphatidylserine (PS) level in the parasite after knock-down of *Pf*LPL1. **(B)** Analysis of major neutral lipids levels showing upregulation of diacylglycerols (DAGs) and downregulation of triacylglycerols (TAGs) in *Pf*LPL1-iKO set. **(C)** Analysis of Fatty Acids (FAs) showing no significant changes in the FA composition of parasites from control and *Pf*LPL1-iKO sets.

### Reduction in *Pf*LPL1 levels disrupts hemozoin formation in the parasite

The localization pattern, knockdown and lipidomic analysis clearly suggested involvement of *Pf*LPL1 in generation/storage of the neutral lipids in the parasite; neutral lipids are known to be efficient catalysts of hemozoin formation in the parasite (11, 26–29). Our data also showed that down-regulation of *Pf*LPL1 caused food-vacuole abnormalities associated with hemozoin development in the parasites. To ascertain the relation between hemozoin development and reduction in neutral lipids after down-regulation of *Pf*LPL1, we assessed total hemozoin content after selective knockdown of *Pf*LPL1 in the transgenic parasites. Total hemozoin content was estimated in equal number of trophozite stage parasites grown in control (with TMP) and *Pf*LPL1-iKO (without TMP) sets. The transgenic parasites growing in *Pf*LPL1-iKO set showed ∼40% reduction in hemozoin levels as compared to control parasite. The wild-type *P. falciparum* 3D7 parasite culture treated with chloroquine, which is known to interfere with hemozoin formation (30, 31), was used as positive control, these parasites showed ∼45% reduction in hemozoin levels (Figure 8). The control parasite line PM1KO, showed no difference in hemozoin levels when grown in presence or absence of TMP.

**Figure 8:**
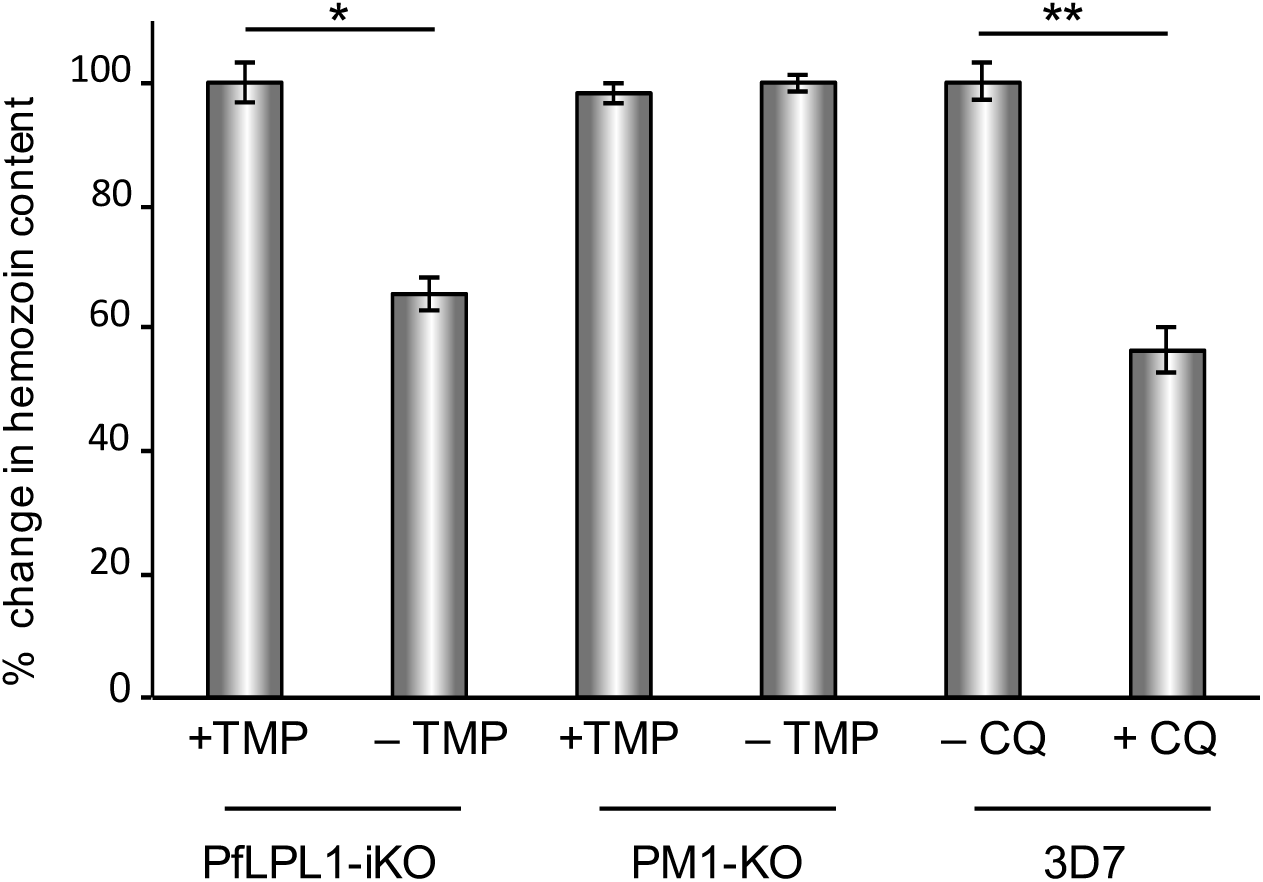
Down-regulation of *Pf*LPL1 protein in the transgenic parasites and its effect on the formation of hemozoin in the parasites. Synchronous transgenic parasites at ring stages were grown till late trophozoite stages and total hemozoin was purified and estimated spectroscopically. Graph showing reduction in hemozoin content in transgenic parasites *Pf*LPL1-iKO (-TMP) as compared to control parasites (+TMP). The parent PM1-KO parasite line was used as control. The 3D7 parasite line treated with chloroquine was used as positive control showing significant reduction as compared to untreated controls. The *p* values were calculated by Student’s t-test: * *p* <0.01 and ** *p* <0.005

## Discussion

The establishment of the intra-erythrocytic *P. falciparum* infection is associated with a large increase in the phospholipid, neutral lipid and lipid-associated fatty acid (FA) content in infected red blood cells (13). In addition to *de novo* synthesis, scavenging of lipids/phospholipids from host milieu and their catabolism may play crucial role to fulfill parasite requirement of lipids during the erythrocytic cycle. In our attempts to further understand the process of catabolism of phospholipids in the parasite, we characterized a putative lysophospholipase in *P. falciparum*, *Pf*LPL1, which is member of the family of lysophospholipase-like protein in the parasite. The *Pf*LPL1 harbours an α, β-hydrolase domain and a putative peptidase domain with conserved catalytic triad. The LPLs are important cellular enzymes that hydrolyze lysophospholipids, an important junction point in the phospholipid metabolism. Indeed, biochemical characterization of recombinant *Pf*LPL1 showed that it harbors lysophopholipase activity. Further, we carried out detailed localization and functional studies to understand significance of the *Pf*LPL1 for blood stage parasite survival. Novel tools have been developed for transient down regulation of proteins in *Plasmodium* by tagging native genes with degradation domain etc. (32–35). Here we have used the regulated fluorescent affinity (RFA) tag system for C-terminal tagging of the native gene, so that the transgenic parasites express *Pf*LPL1 fused with a GFP (or HA) and DHFR degradation domain (DDD or DD) (21).

Confocal microscopic studies using transgenic parasites expressing *Pf*LPL1-RFA fusion protein showed that the protein is associated with a vesicular system in the parasite. The parasite is known to uptake content of host erythrocyte cytosol, especially the host hemoglobin, as a major source of amino acids. The hemoglobin is taken up through the cytosome into double membrane bound vesicles, cytostomal vesicles, which subsequently fuse with the food vacuole (36, 37), during this process the outer membrane of these vesicles fuses with the food-vacuole membrane, whereas the inner membrane is suggested to get degraded by phospholipases (38). However, the *Pf*LPL1 associated vesicles were not found to fuse with the food-vacuole, neither the *Pf*LPL1 staining was seen inside the food-vacuole; rather the *Pf*LPL1-vesicles formed a multi-vesicular structure in close proximity to the food-vacuole. Immuno-electron microscopic studies also showed that the *Pf*LPL1 is enclosed in the membrane bound vesicles near parasite membrane and in the parasite cytosol. Further, localization of fusion protein in large vesicular structures near the food-vacuole, showed resemblance with the previously reported neutral lipid bodies, which are known to be intimately associated with the food vacuole (11). These lipid bodies are the sites of storage of di-and tri-acylglycerols (DAGs and TAGs). These neutral lipids are possibly obtained by the parasite by scavenging or these are generated by digestion of phospholipids during membrane recycling (39, 40). Co-staining of neutral lipid body in the transgenic parasites confirmed association of *Pf*LPL1 with these lipid bodies. These results suggest that the *Pf*LPL1 is involved in generation of neutral lipids from complex phospholipids; these phospholipids may have been taken up by the parasite from host. The malaria parasite is known to uptake exogenous lipids and lipid precursors during blood stage cycle (24, 25). *P. falciparum* blood stages are also able to survive in a minimal medium, only containing two sources of fatty acids, C16:0 and C18:1, suggesting that host scavenging is important for the parasite survival (41, 42). Incorporation of exogenous lysophosphocholine is shown to be necessary for growth of *Plasmodium* (43). It is also shown that *Plasmodium* infected erythrocytes are able to scavenge substantial amount of albumin bound lysophosphocholine from host plasma that is catabolized to generate free fatty acids that are used for *de novo* lipid synthesis in the parasite (44). Furthermore, the apicoplast lipid composition seems to be largely depending on lipid scavenged from the host during blood stages as well (24). The lysophosphatidylcholines (LysoPCs) and phosphatidylcholines (PCs), the major phospholipids detected in the human serums, could be taken up by the parasite (16). It is also shown that phosphocholine can be readily transferred from the erythrocyte membrane to intraerythrocyte parasite suggesting presence of a selective machinery for phosphocholine transport from host cell to the parasite (45). Indeed, a phospholipid transfer protein is identified in *P. falciparum* that can transport different phospholipids between the membranes, this transporter is exported into the erythrocyte cytosol and it is closely associated with parasitophorous vacuole (46). Although, the exact role of this transporter is not clear, but it is shown to be essential parasite survival (47); it may be involved in transport of phosphocholine and other phospholipids from the erythrocyte membrane to the parasitophorous vacuole. These phospholipids may be taken up by the parasite by endocytosis or along with the host cytosol uptake. Our data showed that the *Pf*LPL1 gets associated with small endocytic vesicles near parasitophorous vacuole and neutral lipid stores, which indicate that the *Pf*LPL1 is involved in catabolism of phospholipids/lysophospholipids as these are scavenged from the host or taken up during the process of endocytosis. Furthermore, recent studies showed that the asexual intra-erythrocytic development of *P. falciparum* depends on the scavenging of lysoPC from the extracellular medium to maintain PC synthesis, membrane biogenesis and parasite proliferation (16). The parasite recycles the phosphocholine from lysoPC but not much is known on the molecular mechanism allowing the release of the polar head and the fate of the remaining FA/mono-acylglycerol left after phosphocholine recycling. Such enzymatic activity could well be performed by *Pf*LPL1 in the parasite. The action of *Pf*LPL1 on its substrate, such as lysophosphocholine, will generate fatty acids and glycerophosphocholine, which can be subsequently catabolized by enzyme glycerolphosphodiestere phosphodiesterases (GDPD) to generate glycerol-3-phosphate and choline. The recombinant *Pf*LPL1 was able to carry out this reaction *in vitro*. Further, the GDPD has recently been identified and characterized in *P. falciparum* (48).

The transient down-regulation of *Pf*LPL1 using RFA system (21) caused reduction in neutral lipid stores. This strongly indicates that *Pf*LPL1 is involved in production of neutral lipids in the parasites. TAGs are the major acylglyecrol detected in the infected erythrocytes (13). Further, the Nile Red staining studies suggest that these TAGs could be majorly stored in the lipid storage body associated with the food-vacuole. The TAGs are synthesized through a three sequential acylation steps from a glycerol-3-phosphate backbone. Usually, first two steps are catalyzed by glycerol-3-phosphate acyltransferase (AGPAT), producing LysoPA (LPA). AGAPT are present in two copies of different origins in the parasite genome, one encoding a eukaryotic AGPAT in the ER, PfAGPAT and seems essential for blood stages(49), whilst the other one is of plant origin and present in the apicoplast, *Pf*apiG3PAT or ATS1and is rather essential in liver stages and in *T. gondii* where it generates LPA backbone for the bulk phospholipid synthesis (10, 50, 51). The second step is catalyzed by acyl-glycerol-3-phosphate acyltransferase (AGPAT), which use LPA to add a second FA and form PA. AGPATs are also present in two copies of different origin (eukaryotic and possibly plant-like) and seem important for parasite survival but their exact role for lipid synthesis is yet to be determined. The PA formed by GPATs and AGPATs is then dephosphorylated by phosphatidic acid phosphatases (PAP) to form DAG. Putative PAPs have not been characterized yet and their function and role remain to be elucidated although the parasite genome seems to express 2-3 putative PAPs(52). The final acyl transfer is carried out by the enzyme acyl CoA: DAG acyltransferase (DGAT), which transfer a third fatty acid onto the *sn*-3 position of the glycerol-3-phosphate backbone from DAG. It is shown that *P. falciparum* DGAT activity for *de novo* TAG biosynthesis and encodes a single DGAT homologue highly expressed in late trophozoite and schizont stages that may be essential for parasite survival (13, 39). This suggests that the glycerol-3-phosphate, DAG, and FA that are generated by catabolism of phospholipids by *Pf*LPL1 and *Pf*GDPD, and de novo assembly via GPATs and AGPATs can subsequently be utilized for biosynthesis of TAGs. Therefore, lipid recycling in the parasite involves catabolism of phospholipids and synthesis of neutral lipids simultaneously.

Our lipidomic analysis revealed that there was no change in the FA composition of parasites lacking *Pf*LPL1, suggesting that the enzyme does not have a specific FA species substrate affinity, which reduction(s) would have been quantified. Reduction of PA content in the mutant could indicate that the FA generated via the action of *Pf*LPL1 could fuel the ER resident acyltransferases GPAT and/or AGPAT putatively responsible for the PA *de novo* synthesis (49).PA being used as a precursor for PE synthesis via the Kennedy pathway (6, 53), the reduced PA levels would thus impact PE synthesis. Similarly, PS requires the scavenging of serine, which is massively incorporated for the synthesis of PS via the CDP-DAG pathway. Its synthesis would be favored against the Kennedy pathway due to the lack of its PA substrate. The PC increase might be linked to (i) putative polar head exchange between PS and PC as previously suggested (6), or (ii) the putative increase in lysoPC not being degraded into FA+glycero-3-phosphocholine and thus directly used to form PC by the action of acyltransferases and *Pf*PMT (16, 54) generating phosphocholine from the unused phosphoethanolamine that is normally used for PE synthesis. Importantly, our results clearly point at a significant reduction of TAGs upon *Pf*LPL1 disruption, correlating and confirming our cellular evidences for the role of this enzyme in TAG synthesis, likely by catabolism of scavenged host/external environment phospholipids. The concomitant increase of DAG upon *Pf*LPL1 disruption intuitively suggests that DAG constitutes the likely acceptors of FA provided by *Pf*LPL1 to form TAGs via the action of the sole DGAT of the parasite. Further work would be interesting to determine which is the direct scavenged substrate of *Pf*LPL1, although, phospholipid fluctuations suggest that it could well be the LysoPC, which has been shown to be a key lipid metabolite for asexual development (16, 17). Transient down regulation of *Pf*LPL1 caused growth retardation of transgenic parasites. Further, the parasite developmental-stage profile showed that the development of trophozoites into schizonts was severely disrupted in these parasites. Overall these results showed that *Pf*LPL1 plays functionally important role in parasite growth and development; the *Pf*LPL1 is involved in catabolism of phospholipids which is needed for production of TAGs in the parasite; down-regulation of *Pf*LPL1 and subsequent reduction in TAGs stores caused severe growth retardation in the asexual stage parasites.

The exact role of TAGs stores in the parasite is not very clear, however, our results and previous studies with lipid labeling dyes showed that there is active synthesis of TAGs in the asexual stage parasite (11). It is suggested that TAGs stores may not be involved *de novo* synthesis of membrane lipids and membrane biogenesis (39). Further the parasite may not be able to use TAGs as substrates for oxidative catabolism as unlike *T. gondii* it lacks most enzymes involved in beta-oxidation of FA (55). Our data shows that *de novo* synthesized TAGs in the parasite might play important role its growth and development. The close association of TAGs stores with the food-vacuole suggest that their role in parasite is associated with processes in the food-vacuole. The parasite food-vacuole is mainly involved in proteolytic degradation of host hemoglobin and sequestration of released haem in the form of hemozoin; the haem crystallization pathways is the major target of known drugs and is also targeted to develop new anti-malarials (56). Several lines of studies have suggested that neutral lipids might play important catalytic role in hemozoin formation; the neutral lipids form nanosphere provide unique environment that promote haem polymerization (11, 26–29). Our studies showed that down-regulation of *Pf*LPL1 and subsequent reduction in TAGs stores effected development of parasite at trophozoite stages with abnormal food-vacuole morphology without clearly developed hemozoin crystal. The total hemozoin estimation data correlates with these observations, as there was clear reduction in hemozoin levels after *Pf*LPL1 knock-down in the parasites. These results suggest that TAGs stored in close association with the parasite food-vacuole are involved in hemozoin development.

Overall our data showed that down-regulation of *Pf*LPL1 in the parasite reduced *de novo* synthesis of TAGs; reduction in TAGs stores hampered hemozoin development in the food-vacuole, which subsequently caused reduction in parasite growth, and development. The metabolic pathways of phospholipid recycling and *de novo* lipid synthesis are therefore crucial for parasite survival. Detailed understanding of neutral lipid-based heme detoxification via TAG and inhibition of *Pf*LPL1may help us to design efficient novel anti-malarial strategies.

## Methods

### Parasite culture, plasmid construct and parasite transfection

*Plasmodium falciparum*strain 3D7 was cultured in RPMI media (Invitrogen) supplemented with 0.5% albumin4% haematocrit and parasite cultures were synchronized by repeated sorbitol treatment. To generate a transfection vector construct, a C-terminal fragment of *pfLPL1*gene (241-1059 bp) was amplified from *P. falciparum*3D7 genomic DNA using primers 971A (5’-*CTCGAG*ATATATGAAGGTAGTTGGATTG-3’) and 972A (5’*CCTAGG*TTCACATTTTTTTATCCAAG-3′). The amplified PCR product was digested with *Xho*I and *Avr*II restriction enzymes and cloned in frame to the N-terminus of GFP in the *Xho*I and *Avr*II sites of the vector pGDB (21) to yield construct pGDB-*Pf*LPL1. The plasmids were transfected into *P. falciparum* Plasmepsin-I knockout parasite line (PM1KO),which contains the human DHFR (hDHFR) integrated into a non-essential gene(57). Synchronized ring-stage parasites were transfected with 100 μg of purified plasmid DNA (PlasmidMidi Kit, Qiagen, Valencia, CA) by electroporation (310 V, 950 μF) (58).Transfected parasites were selected over 2.5 µg/ml blasticidin (Calbiochem) and 5 µM trimethoprim (TMP) (Sigma) and subsequently subjected to on and off cycling of blasticidin drug to promote integration of the plasmid in the main genome. Integration was assessed by PCR based analysis and parasites with integration were obtained after two rounds of blasticidincycling. The TMP was always present in the medium after its initial introduction.

### Isolation of parasites and Western immunoblotting

For Western blot analyses, parasites were isolated from tightly synchronized cultures at trophozoite stage by lyses of infected erythrocyte with 0.15% saponin. Parasite pellets were washed with PBS, suspended in Laemmli buffer, boiled, centrifuged, and the supernatant obtained was resolved on 12% SDS-PAGE. The fractionated proteins were transferred from the gel onto a PVDF membrane (Amersham) and the membrane was blocked in blocking buffer (1×PBS, 0.1% Tween-20, 5% milk powder) for 2 h. The blot was washed and incubated for 1h with primary antibody [mice anti-GFP (1:1000); rabbit anti-Bip (1:2000), rabbitanti-SERA (1:2000)] diluted in dilution buffer (1× PBS, 0.1% Tween-20 and 1% milk powder). Later, the blot was washed and incubated for 1 h with appropriate secondary antibody (anti-rabbit or anti-mouse, 1:2000) conjugated to HRP, diluted in dilution buffer. Bands were visualized by using HRP-DAB.

### Fluorescence microscopy and indirect immunofluorescence assay

*P. falciparum* culture transfected with pGDB-*Pf*LPL1 (labelled as *Pf*LPL1-RFA) was synchronized by two consecutive sorbitol treatments 4h apart. Parasites at different developmental stages were collected from the culture for fluorescence microscopy and stained with DAPI at a final concentration of 2 μg/ml for 30 min at 37°C prior to imaging. Fluorescence from DAPI and GFP was observed and captured from live cells within 30 min of mounting the under a cover slip on a glass slide, using Nikon A1 confocal laser scanning microscope.

To visualize the endoplasmic reticulum, the transgenic parasites were stained with ER-Tracker Red CMXRos (Invitrogen) at a final concentration of 20 nM in 1× PBS for 15 min at 37°C. The membrane structure in parasitized erythrocytes were labelled with BODIPY-TR-ceramide (Molecular Probe)(37, 59). Briefly, parasitized erythrocytes were resuspended in complete media (5% parasitemia, 4% hematocrit) and incubatedwith 1µM BODIPY-TR-ceramide (Invitrogen) at 37°C for 60 min, washed in complete media three times and examined by fluorescence microscopy. To label the neutral lipid storage structures, infected erythrocytes were labelled with Nile Red (Molecular Probes) using a modified method of Palacpac *et al.* (60). Briefly, Nile Red was added to a final concentration of 1 µg/ml into the parasite culture (5-10% parasitaemia, 3% haematocrit), the cultures was incubated on ice for 30 min and subsequently washed with 1×PBS before analysis by confocal microscopy.

Indirect Immunofluorescence assays were performed on *P. falciparum*3D7 or transgenic parasite lines as described earlier (61). Briefly, the parasite samples were fixed with 4% paraformaldehyde, incubated with primary antibody (rabbit anti-SERA diluted 1:500 in 3% BSA, 1× PBS) and subsequently with Alexa 594 linked goat anti-rabbit antibodies (1:250, Life Technologies, USA) as secondary antibody with intermittent washing. The parasite nuclei were stained with DAPI (2 μg/ ml).

The GFP expressing parasites as well as the parasite stained with different labelling dyes were viewed using a Nikon A1 confocal laser scanning microscope. For live cells, observations were limited to 30 min to ensure parasite viability throughout the analyses. The 3D images were constructed by using series of Z-stack images using IMARIS 7.0 (Bitplane Scientific) software.

### Cryo-immunoelectron microscopy

Immunoelectron microscopy was carried out on transgenic *P. falciparum* parasites, *Pf*LPL1-RFA, expressing *Pf*LPL1 fused with GFP tag. The trophozoite stage parasites were fixed in 4% paraformaldehyde, 0.04% glutaraldehyde in 1× PBS at 4°C for 1 h and subsequently embedded in gelatin, and infiltrated with a cryo-preservative and plasticizer (2.3 M sucrose/20% polyvinyl pyrrolidone). After freezing in liquid nitrogen, samples are sectioned with a Leica Ultracut UCT cryo-ultramicrotome (Leica Microsystems, Bannockburn, IL) at −260°C. Ultra-thin sections were blocked with 5% fetal bovine serum and 5% normal goat serum in 1× PBS for 30 min and subsequently stained with rabbit anti-GFP antibody (Abcam, 1:500 dilution in blocking buffer), washed thoroughly and incubated with 18 nm colloidal gold-conjugated anti-rabbit IgG for 1 h. Sections were stained with 0.3% uranyl acetate/1.7% methyl cellulose and visualized under a JEOL 1200EX transmission electron microscope (JEOL USA, Peabody, MA). All labelling experiments were conducted in parallel with controls omitting the primary antibody or using pre-immune sera as primary antibodies.

### *In vitro* parasite growth inhibition assay

For growth assay tightly synchronized parasites at ring stage were cultured in a 6 well plate. The assay was performed in triplicate. Each well was containing 4% haematocrit, 1ml of complete media supplemented with 5µM TMP and the parasitaemia was adjusted to 0.5%. A parallel set of parasite culture was grown in media without TMP. For microscopic analysis, smears were made from each well at different time points (0, 24, 48, 72, 96 and 120 h) stained with Giemsa, and the numbers of ring/trophozoite stage parasites per 5000 RBCs were determined and percentage parasitemia was calculated to assess the parasite inhibition.

### Lipid isolation and mass spectrometry-based lipidomic analyses

Infected red blood cells (5×10^7^ cell equivalents) were harvested from control *Pf*LPL1-iKO cultures and parasites released from the erythrocyte hosts by saponin treatment as previously described (24). The parasite pellets were washed in PBS and stored at −80°C or directly used for further analysis. Their total lipid spiked with 20 nmol C21:0 phosphatidylcholine was extracted by chloroform: methanol, 1:2 (v/v) and chloroform:methanol, 2:1 (v/v). The pooled organic phase was subjected to biphasic separation by adding 0.1% KCl and was then dried under N_2_ gas flux prior to being dissolved in 1-butanol. For the total fatty acid analysis, an aliquot of the lipid extract was derivatized on-line using MethPrep II (Alltech) and the resulting FA methyl esters were analyzed by GC-MS as previously described(10). For the quantification of each lipid, total lipid was separated by 2D HPTLC using chloroform/methanol/28% NH_4_OH, 60:35:8 (v/v/v) as the 1st dimension solvent system and chloroform/acetone/methanol/acetic acid/water, 50:20:10:13:5 (v/v/v/v/v) as the 2nd dimension solvent system. Each lipid spot was extracted for quantification of fatty acids by gas chromatography-mass spectrometry (Agilent 5977A-7890B) after methanol lysis. Fatty acid methyl esters were identified by their mass spectrum and retention time, and quantified by Mass Hunter Quantification Software (Agilent) and the calibration curve generated with fatty acid methyl esters standards mix (Sigma CRM47885). Then each lipid content was normalized according to the parasite cell number and a C21:0 internal standard (Avanti Polar lipids). All analyses were performed in triplicate or more (*n*=3). * P values of ≤ 0.05 from statistical analyses (Student’s *t*-test) obtained from GraphPad software analysis were considered statistically significant.

### Hemozoin isolation and estimation

Total hemozoin was purified following Coban *et al.*(2002)(62). Briefly, equal number of parasitized erythorcytes with trophozite stage parasites were collected from synchronize *P. falciparum* cultures. Parasites were harvested by saponin lysis, washed five times with phosphate-buffered saline (PBS) and sonicated in 2% SDS. The pellet was washed seven to eight times in 2% SDS, resuspended in of 10 mM Tris-HCl (pH 8.0), 0.5% SDS, and 1 mM CaCl_2_ containing proteinase-K (2 mg/ml) and incubated at 37°C overnight. The pellet was again washed three times in 2% SDS and incubated in 6 M urea for 3 h at room temperature with shaking. The pellet was again washed three times in 2% SDS and then in distilled water. The final hemozoin pellet was resuspended in distilled water. Total heme content was determined following Tripathi *et al*. (63). The hemozoin polymer was depolymerised by incubation in 1 ml 20mM NaOH/2% SDS at room temp for 2 h. The absorbance of solution was taken at 405 nm and heme concentration was calculated from standard curve. Heme stock (10mM) was prepared by dissolving 3.3mg of hemin (Sigma) in 500 µl of 1 M NaOH, which was used to make dilutions ranging from 50µm-600µm to plot the standard curve.

### Flow cytometry

The change in levels of GFP tagged protein and neutral lipids stored in transgenic parasites were assessed by flow cytometry. Tightly synchronized ring stages transgenic parasite cultures were grown in media containing 5μM TMP or solvent alone, aliquot of the parasite samples were collected after 24h. GFP fluorescence and Nile Red staining was quantified using the BD FACS Calibur system (Beckton Dickinson). Arbitrary fluorescence units for 100,000 cells were acquired on channels FL1 (GFP) and FL2 (Nile red) using CellQuest Pro (Beckton Dickinson) and data were analyzed using Graph Pad Prism v 5.0. Uninfected RBCs were used as background control.

### Statistical Analysis

The data sets were analyses and graphical presentations were made using GraphPad Prism ver 5.0, and the data were compared using unpaired Student’s *t*-test.

## Acknowledgements

We are grateful to Daniel Goldberg and Vasant Muralidharan for pGDB vector and PM1-KO parasite line. We thank Wandy Beatty for help with the immunoelectron microscopic studies, and Rotary blood bank, New Delhi, for providing the RBCs. MA is supported by research fellowship from ICMR, Govt. of India. VT is supported by BioCARE grant from Department of Biotechnology, Govt. of India.CYB and YYB are supported by Agence Nationale de la Recherche, France (Grant ANR-12-PDOC-0028-Project Apicolipid), the Atip-Avenir and Finovi programs (CNRS-INSERM-FinoviAtip-AvenirApicolipidprojects), and the Laboratoire d’ Excellence Parafrap, France (grant number ANR-11-LABX-0024). The research work in AM’s laboratory is supported by Centre Of Excellence grant (BT/COE/34/SP15138/2015) from Department of Biotechnology, Govt. of India. The research work is supported by the CEFIPRA Collaborative Research Program Grant (Project 6003-1) to CYB and AM laboratories.

## Supplementary data

### Identification and sequence analysis of *P. falciparum* LPL1 homologue (*Pf*LPL1)

To determine enzymes responsible for FA trafficking from the host, *in silico* analyses was carried out to identify *P. falciparum* lyso-phospholipases from genome database. *P. falciparum* genome harbors 13 putative lyso-phospholipases and show high degree of homology among themselves. They are around ∼400 amino acid long (except 2 which are a bit longer) that harbor putative lysophospholipase domain (Hydrolase_4 domain; Pfam 12146) within the alpha-beta hydrolase fold (*αβ−*hydrolase_1, Pfam00561). The hydrolase domain is about ∼200-250 amino acid in length. Two members of this *P. falciparum* lyso-phospholipase family, PF3D7_1476700 and PF3D7_1476800, also harbor a peptidase domain (Peptidase_S9, Pfam 00326) belonging to Prolyl oligopeptidase family. SSDB motif search indicates the presence of peptidase domain in a couple of other lysophospholipase but is not considered because of its low E value score. The characteristic GXSXG motif embedding the catalytic active S as well as the aspartate (D) and histidine (H) residue forming the catalytic triad, were found to well conserved in all the lysopospholipases (1, 2). In the present study, we have characterized PF3D7_1476700 (*pfLPL1*). The PfLPL1 is 353aa long protein that harbors a central lysophospholipase domain (27-111 aa; E value 3.97 e^-10^) and a Peptidase_S9 domain (265-352aa; 3.29 e^-6^) (Table S1). Homologues of PfLPL1 are also identified from *P. berghei* (PBANKA_122030), *P. chabudi chabudi* (PCHAS_13706), *P. vivax* strain Sal-1 (PVX_096960), *P. knowlesi* (PKH_010790) and *P. cynomolgi* (PCYB _104230) using the genome database. An alignment of the predicted proteins sequences of these genes showed that the *Pf*LPL1 is highly conserved among these *Plasmodium* species (Figure S1). A structural signature of lipase enzyme is possession of the catalytic triad which is important for its catalytic activity; sequence alignment found Serine-167, Aspartate-300 and Histidine-330 residues to be involved in the formation of the catalytic triad, where the Histidine-330 residue is spatially positioned between the Serine-167 and the Aspartate-300. The nucleophilic serine residue of the active center is present in a penta-peptide sequence GYSMG which is similar to the classic lipase motif GXSXG of α/β hydrolase (Figure S1 and S2).

### Endogenous tagging of *pfLPL1* gene with DD-tag, expression and localization of fusion protein in the transgenic parasites

To generate a transfection vector construct, a C-terminal fragment of *pfLPL1* gene (241-1059 bp) was cloned in *Xho*I and *Avr*II restriction sites of the vector pHADD in frame with HA-DD cassette to yield construct pHADD-LPL1 (Figure S5). The plasmids were transfected into *P. falciparum* as described in method section. Transfected parasites were selected over 2.5nM WR99210 drug and 1μM Shield1; selected parasites subsequently subjected to on and off cycling of WR99210 drug to promote integration of the plasmid in the main genome. Integration was assessed by PCR based analysis and parasites with integration were obtained after three rounds of WR99210 cycling (Figure S5B). The Shield1was always present in the medium after its initial introduction. Western blot analysis confirmed expression of fusion protein, *Pf*LPL1-HA-DD, in the transgenic parasites (Figure S5C). Further immune-staining with anti-HA antibody showed localization of fusion protein in vesicular structures in parasite cytosol (Figure S6A, B); these vesicles merged with a large multi-vesicular like structure near food-vacuole which overlapped with Nile-red staining (Figure S6B). These results confirmed the localization studies with HA-tagged native protein are similar as the RFA-tagged protein in the transgenic parasites.

### Immuno localization assay

Indirect Immunofluorescence assays were performed on *P. falciparum* 3D7 or transgenic parasite lines as described earlier (3). Briefly, the parasite samples were fixed with 4% paraformaldehyde, incubated with primary antibody [rabbit anti-HA (1:1000 diluted in 3% BSA, 1× PBS) and subsequently with Alexa 594 linked goat anti-rabbit antibodies (1:250, Life Technologies, USA) as secondary antibody with intermittent washing. The parasite nuclei were stained with DAPI (2 μg/ml). The parasite were observed under the confocal laser scanning microscope.

### Knock-down of *Pf*LPL1 by selective degradation through DD-tag, its effect on parasite growth and neutral lipids

Selective degradation of DD-tagged protein in absence of Shield-1was assessed by western blot analysis; as shown in Figure S7A, removal of Shield-1 resulted in down regulation of *Pf*LPL1-HA. Further, effect of selective degradation of DD-tagged protein on parasite growth was estimated by counting new ring stage parasites in subsequent cycles. As shown in Figure S7B, in absence of Shield-1 (*Pf*LPL1-iKO), the parasite growth was significantly reduced (∼80%) as compared to control parasite culture (+Shield-1). To confirm the effect of down regulation of DD tagged *Pf*LPL1 on levels of neutral lipid in lipid storage body, synchronized ring stage *Pf*LPL1-DD parasites were grown in presence or absence of Shield-1 for 24-30h to develop into trophozoite stages, subsequently levels of neutral lipids were assessed in these parasites by Nile Red staining. Downregulation of *Pf*LPL1 resulted in loss in neutral lipid levels in the lipid storage body as compared to control parasites (Figure S7C).

### Expression, purification of recombinant *Pf*LPL1 and enzyme activity assay

In view to characterize the biochemical activity of *Pf*LPL1, recombinant protein corresponding to the full length *pfLPL1* gene was expressed using *E. coli* expression system. A DNA fragment corresponding to full length *pfLPL1* gene (1aa-353 aa) containing the lipase domain was amplified by PCR using *P. falciparum* 3D7 genomic DNA with primers 966A and 967A. The amplified fragment was cloned in the *Nco*I and *Xho*I sites of pETM-41 expression vector. The resultant plasmid pETM-41-*pfLPL1* was transformed into *E. coli* BL21(DE3) cells for expression of the recombinant protein. These *E. coli* BL21(DE3) cells were grown in Luria broth containing Ampicillin (100µg/ml) at 37°C under shaking to an OD_600_ of 0.6-0.7 and expression of recombinant protein was induced with isopropyl-*β*-thioglactopyranoside (IPTG) at a final concentration of 1 mM. The recombinant *Pf*LPL1 (∼73 kDa) abundantly expressed as a soluble protein in *E. coli* cells and was purified by two step affinity chromatography using Ni^2+^ NTA and amylose resins (Figure S8 A & B).

The purified recombinant PfLysoPL1 was assessed for its lysophospholipase activity using an *in vitro* lysophospholipase assay modified from Kishimoto *et.al* (4) using Amplex® Red Phospholipase kit as per manufacturer’s instructions (5). The reaction mechanism of the enzymatic assay involves two step hydrolysis of LPC by lysophospholipase to glycerophosphoryl-choline (GPC), which is further hydrolysed to generate glycerophosphate and choline. The choline is further oxidized by choline oxidase to betaine and H_2_O_2_. Finally, H_2_O_2_, in the presence of horseradish peroxidase, reacts with Amplex Red reagent (10-acetyl-3,7-dihydrophenoxazine) to yield a highly fluorescent product, resorufin, fluorescence in the reaction mix is measured at 530/590 nm. The assay was performed in a 200 µl volume containing 10µg of recombinant protein, in 1X reaction buffer (250 mM Tris-HCl, 25mM CaCl_2_, pH 8.0), 20mM H_2_O_2_, 100µM Amplex Red reagent, 2 U/mL HRP, 0.2 U/mL choline oxidase, 0.5mM lecithin and 0.1U GPCP. The enzyme activity is recorded as the increase in fluorescence (excitation 530 nm; emission 590 nm) for 3-6h at room-temperature using a LS50B Fluorometer (Perkin-Elmer). The standard curve was prepared using different concentration of resorufin and activity of recombinant protein was calculated as a function of generated resorufrin. For calculation of kinetic constants, reactions were set up with varying concentration of the substrate (0-70 µM). As shown in figure S8C, the recombinant *Pf*LPL1 showed concentration dependent lysophospholipase activity, *K*m and *V*max were determined using the Graph Pad Prism software (*K*m = 61.2 ± 5.9 µM, *V*max= 2116.67 µM/mg/min).

---

**Table S1:** List of putative phospholipases in *P. falciparum*, their expression pattern in different parasite stages and domain architecture.

| PlasmoDB Gene ID | Expression in parasite stages* | Functional Domains by PFAM/SSDB Motif Search (Functional annotation) | Domain length (aa) | Pfam E-value | Protein name (Reference or this study) |
| --- | --- | --- | --- | --- | --- |
| PF3D7_1476700 | R, Sch, G | $\alpha/\beta$ hydrolase, (putative lysophospholipase)<br>Prolyl oligopeptidase | 86-337<br>259-352 | 2.8e-28<br>6e-06 | PfLPL1 |
| PF3D7_1038900 | G-V, Oo | $\alpha/\beta$ hydrolase, putative esterase | 85-340 | 8.1e-29 | PfLPL2 |
| PF3D7_1476800 | R to Sch, G | $\alpha/\beta$ hydrolase, putative lysophospholipase<br>Prolyl oligopeptidase | 86-337<br>256-351 | 2.4e-29<br>7.1e-05 | PfLPL3 |
| PF3D7_0731800 | R, G, Oo | $\alpha/\beta$ hydrolase | 290-503 | 2.6e-22 | PfLPL4 |
| PF3D7_0936700 | R, Sch | $\alpha/\beta$ hydrolase, putative lysophospholipase | 90-141,<br>226-405 | 6.4e-09<br>3.5e-14 | PfLPL10 |
| PF3D7_0702200 | R & Sch, G-V | $\alpha/\beta$ hydrolase, putative lysophospholipase | 86-234,<br>206-385 | 1.3e-08<br>2e-17 | PfLPL20 |
| PF3D7_1252600 | R to Sch | $\alpha/\beta$ hydrolase, putative lysophospholipase | 90-140,<br>171-347 | 2.4e-07<br>6.2e-16 | PfLPL30 |
| PF3D7_1401500 | R, T, G | $\alpha/\beta$ hydrolase, putative lysophospholipase | 90-140,<br>153-345 | 2.5e-08<br>3.4e-13 | PfLPL40 |
| PF3D7_0102400 | G-V | $\alpha/\beta$ hydrolase, putative lysophospholipase,<br>pseudogene | 43-350 | 6.38e-30 | PfLPL50 |
| PF3D7_0709700 | G-V | $\alpha/\beta$ hydrolase, putative lysophospholipase | 95-344 | 7.1e-28 | PfLPL60 |
| PF3D7_0937200 | R, G | $\alpha/\beta$ hydrolase, putative lysophospholipase | 85-338 | 4.6e-28 | PfLPL70 |
| PF3D7_1001400 | R | $\alpha/\beta$ hydrolase, exported lipase 1, lysophospholipase/serine aminopeptidase | 550-767 | 5.7e-18 | PfXL1<sup>#</sup> |
| PF3D7_1001600 | T-Sch | $\alpha/\beta$ hydrolase,<br>(exported lipase 2,<br>lysophospholipase/serine aminopeptidase) | 390-527 | 4.56e-17 | PfXL2<sup>#</sup> |
| *R-ring, T-trophozoite, Sch-schizont, G-gametocyte, Oo-ookinete stages |  |  | #(Spillman *et al.*, 2016) |  |  |

## Supplementary Figures

**Figure S1:**
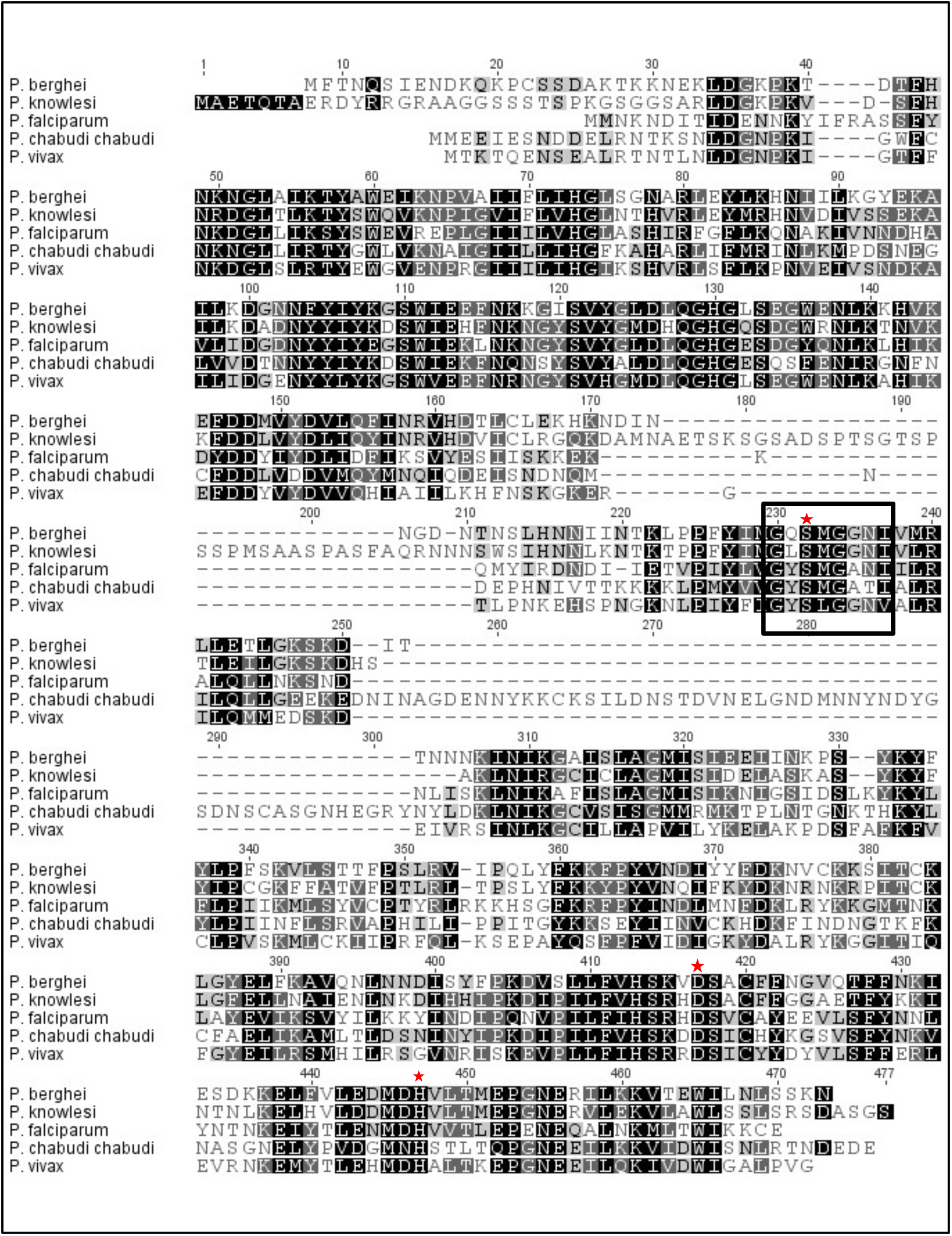
Clustal W alignment of amino acid sequences of lysophospholipase (LPL1) homologues from different species of *Plasmodium*. Amino acids with 100% identity are shown in black, ≥80% in dark grey and ≥60% in light grey. The conserved GXSXG motif is boxed and the conserved catalytic residues (S, D and H) are marked with asterisks.

**Figure S2:**
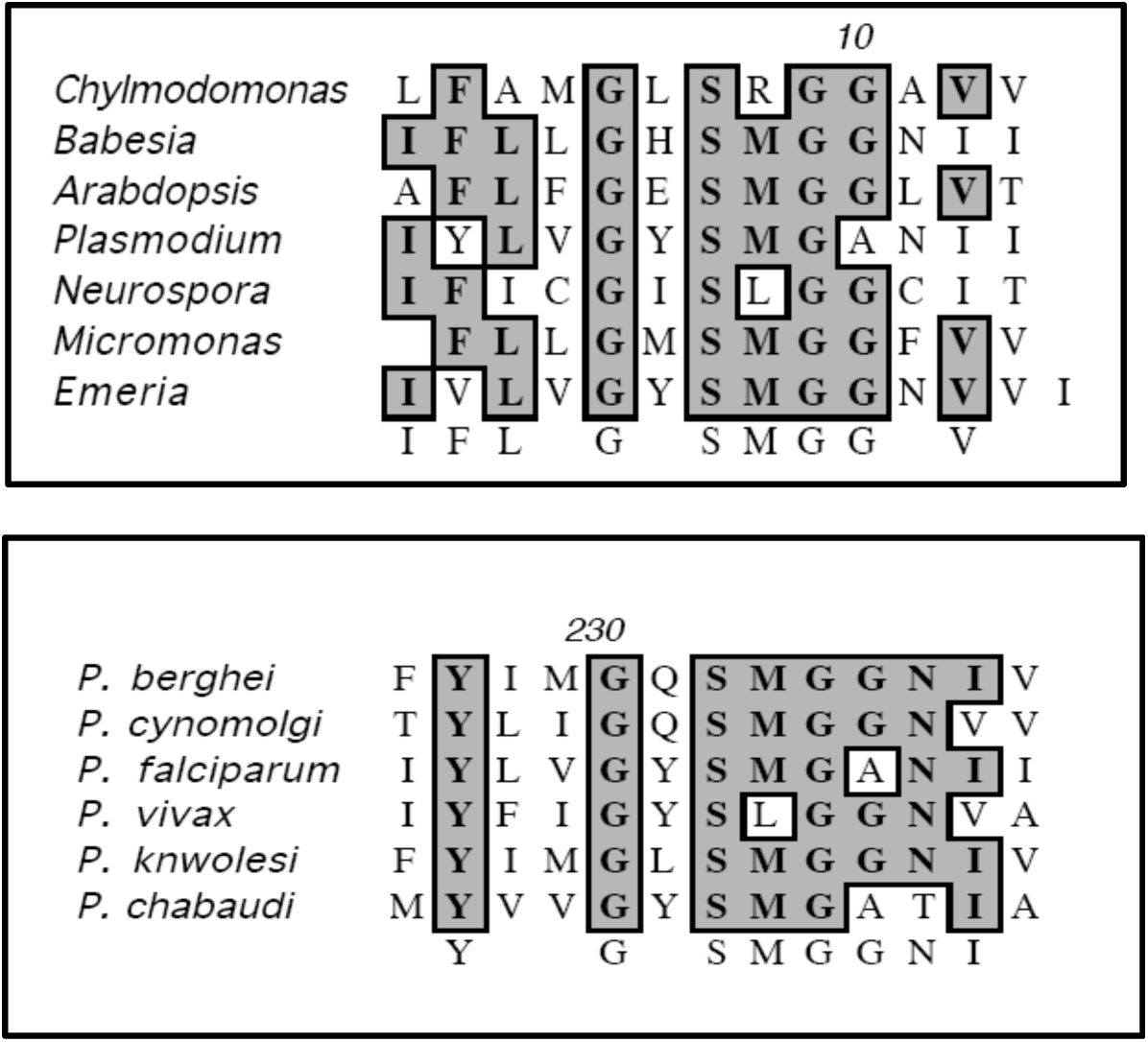
Clustal W alignment of showing the conserved GXSXG motif in homologues of LPL1 of *Emeria tenella* (AET50683.1)*, Micromonas sp. RCC299* (XP_002500615.1)*, Neurospora caninum liverpool (XP_003881427.1), Arabidopsis thaliana (NP_175685.1), Babesia equi* (XP_004833376.1)*, Chylmodomonas reinhardtii* (XP_001690806.1) *and Plasmodium species*. Amino acids with ≥60% identity are shown in grey.

**Figure S3:**
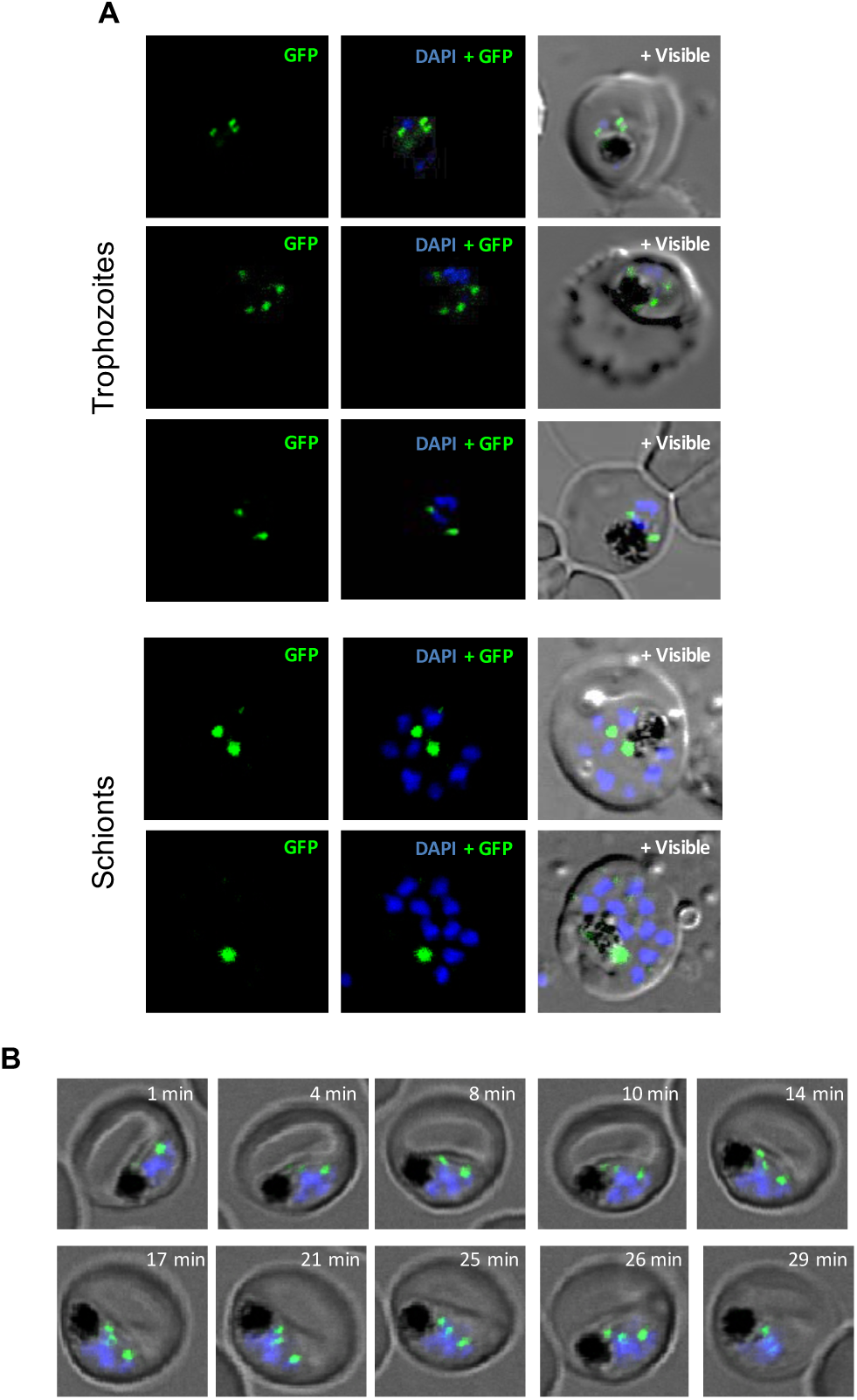
**(A)** Localization of *Pf*LPL1-RFA fusion protein in transgenic *P. falciparum* parasites. Fluorescent microscopic images of live transgenic parasites trophozoite, and stages. The parasite nuclei were stained with DAPI and slides were visualized by confocal laser scanning microscope. The GFP-fluorescence was observed in small vesicles present near the parasitophorous vacuole and in the parasite cytosol. In late trophozoite stage GFP-fluorescence was observed in large vesicular structure in close association with the food-vacuole. **(B)** Consecutive images from time-lapse microscopy of *Pf*LPL1-RFA expressing transgenic parasites showing localization and migration GFP labeled vesicles. Sequential images of a parasite over a time interval of 29 min showing development of a GFP labeled vesicles at the parasite plasma membrane, and series of vesicles traversing in parasite cytosol towards food-vacuole (labelled by presence of hemozoin crystals).

**Figure S4:**
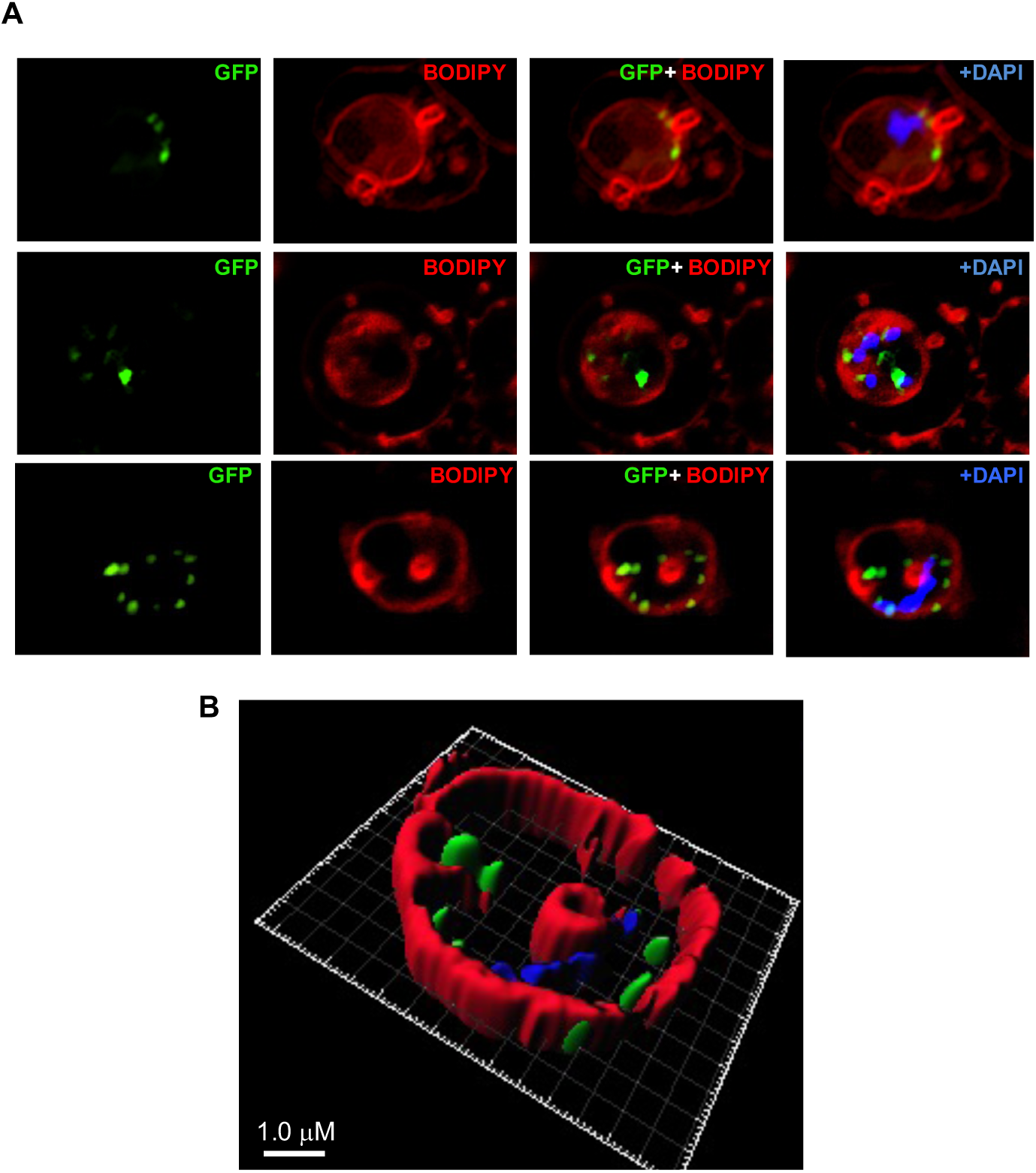
Structured illumination microscopy (SIM) images of live transgenic parasites showing labelling of membranes in infected RBCs and localization of *Pf*LPL1. **(A)** Trophozoite stage transgenic parasites expressing *Pf*LPL1-RFA were stained with BODIPY-TR ceramide (red) and parasite nuclei were stained with DAPI (blue). Small GFP foci of the *Pf*LPL1-RFA fusion protein were observed near parasite boundary (panel 1, marked with arrowhead), these foci showed closed association with fluorescence by BODIPY-TR labelled parasite membrane. In some parasites, the GFP vesicles are seen in parasite cytosol (panel 2 and 3). **(B)** A three-dimensional reconstruction of series of Z-stack images (corresponding to panel 3 in A) using IMARIS software. Small GFP vesicles are present juxtaposed to the parasite membrane and in parasite cytosol.

**Figure S5:**
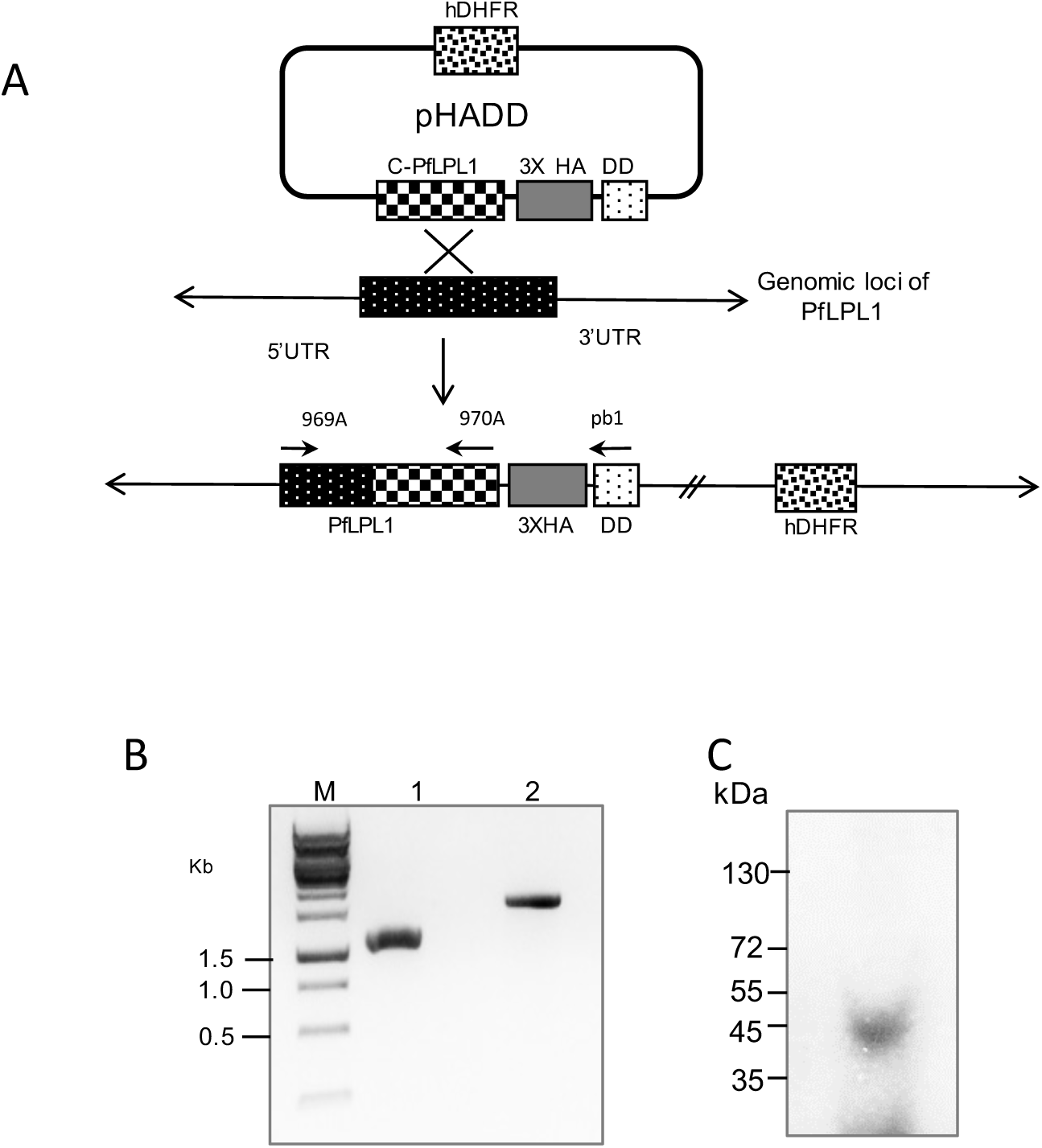
Generation and analysis of *Pf*LPLl-DD parasite line. **(A)** Schematic of pHADD vector illustrating the targeting plasmid for integration in the *Pf*LPLl locus. A fragment corresponding to the 3’ end of the *pfLPL1* gene was cloned upstream of 3XHA tag and ddFKBP gene in the pHADD vector. The primer combinations used PCR analysis to confirm the integration are indicated. **(B)** PCR amplification using genomic DNA from the *Pf*LPLl-DD parasite lines to confirm the integration of the plasmid in the main genome locus. using the primer combination 969A/970A and 969A/Pb1. **(C)** Immunoblot analysis of trophozoite stage *Pf*LPLl-DD transgenic parasites grown in presence of Shld1 drug, using anti-HA antibody. A band of ∼ 53 kDa, representing the fusion protein, is detected in transgenic parasites.

**Figure S6:**
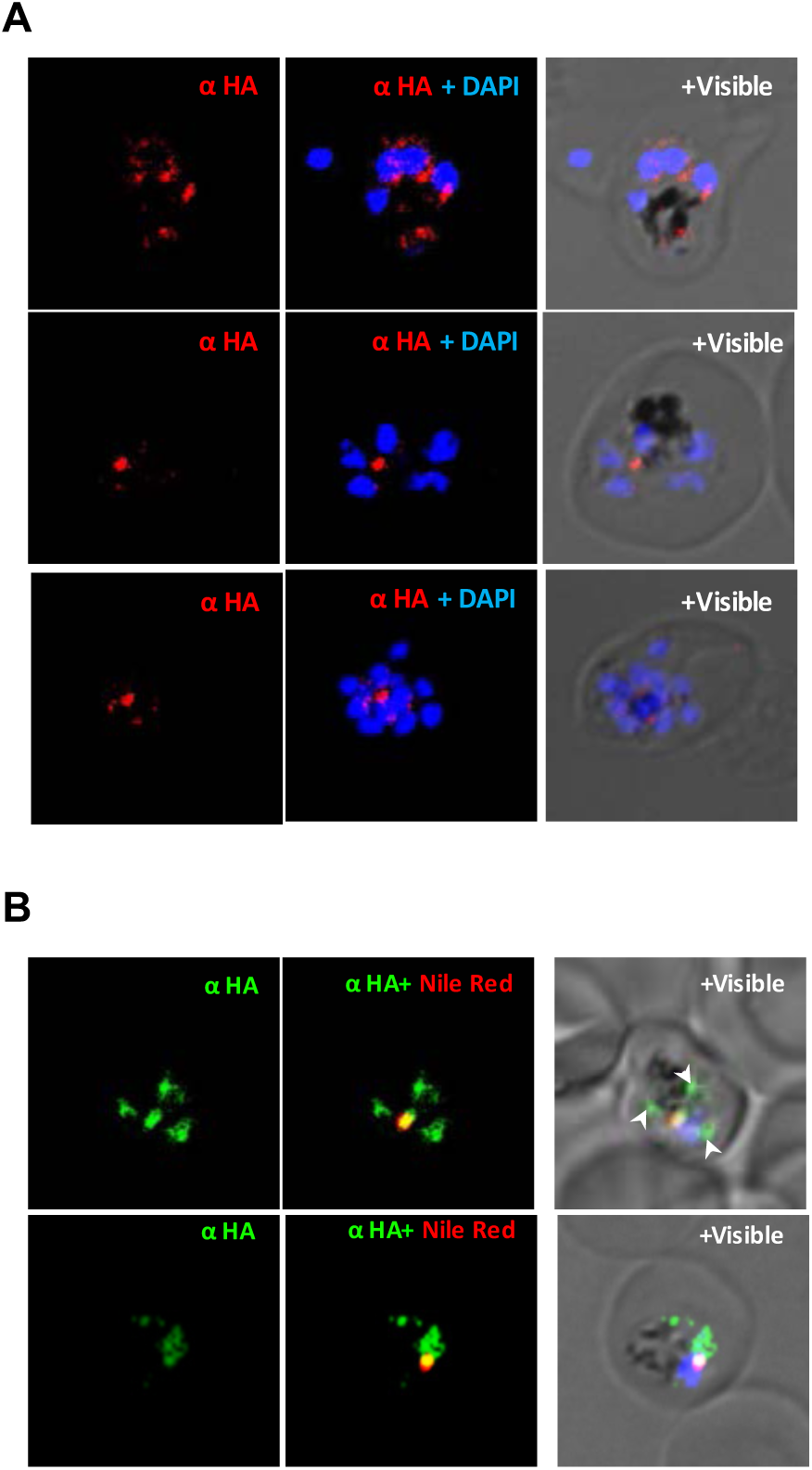
Expression and localization of the *Pf*LPLl fusion protein with ddFKBP degradation domain (*Pf*LPLl-DD) in transgenic parasites. Fluorescence microscopy images of *Pf*LPLl-DD transgenic parasite immuno-stained with anti-HA antibody **(A)**; immuno-stained with anti-PfLPLl antibody **(B)**; and immuno-stained with anti-*Pf*LPLl antibody and co-stained with Nile red **(C)**. The parasite nuclei were stained with DAPI and slides were visualized by confocal microscope. The PfLPL1 was localized in vesicular structures which overlapped in neutral lipid body in late trophozoite stages

**Figure S7:**
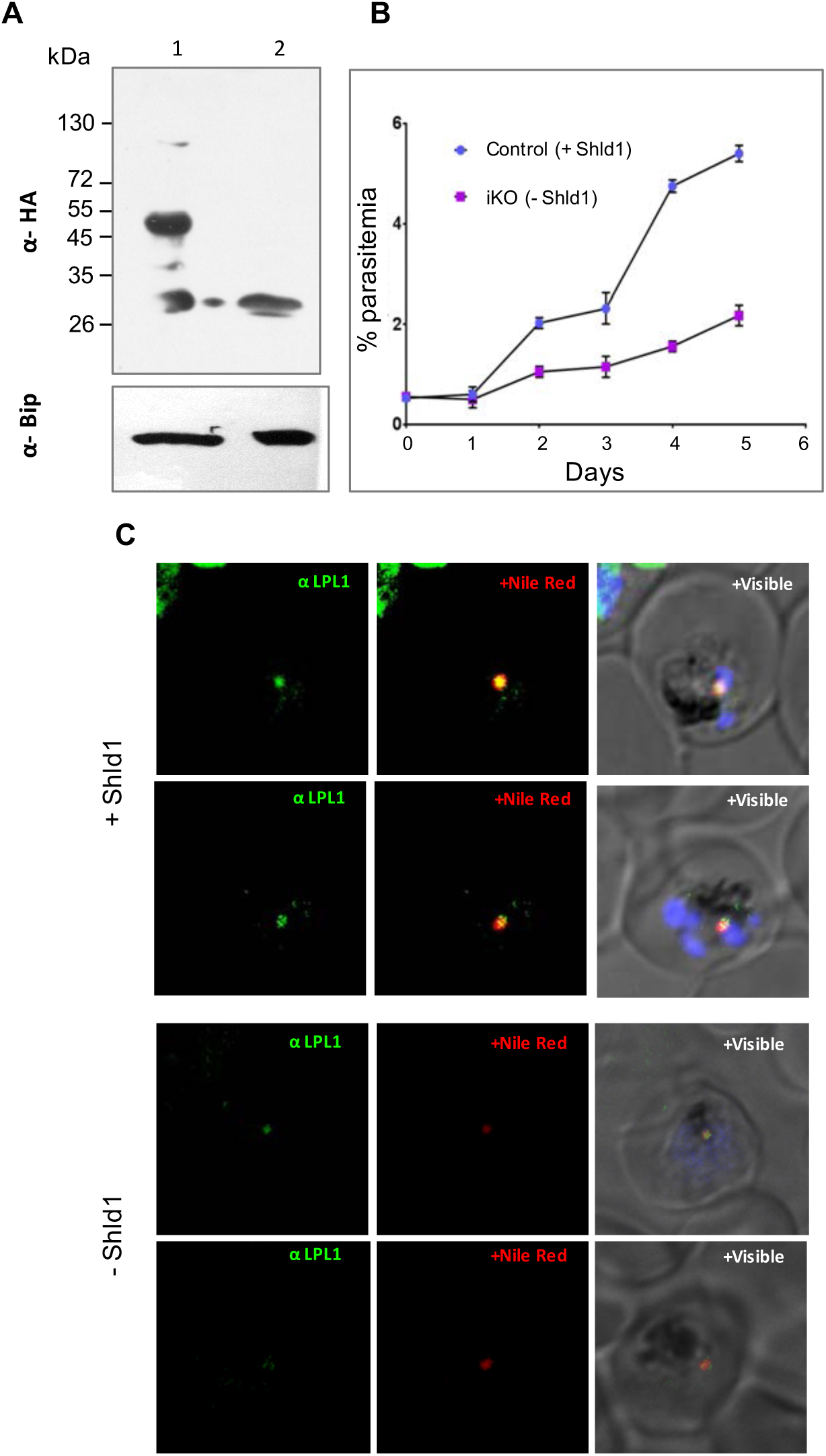
Inducible knock-down of *Pf*LPL1 protein in the *Pf*LPL1-DD transgenic parasites and its effect on growth and development of the parasites. **(A)** Immunoblot analysis using anti-HA antibodies and trophozoite stage transgenic parasites expressing *Pf*LPL1-DD grown in presence of Shld1 drug or solvent alone (control or iKO respectively). A band of ∼ 53 kDa, representing the fusion protein, is detected in the control parasites (lane 1), but not in the *Pf*LPL1-iKO set (lane 2). Parallel blot was probed with anti-Bip antibodies to show equal loading. **(B)** Tightly synchronized ring stage parasite culture (0.2% parasitemia) of transgenic parasites were grown with or without Shld1 (control and *Pf*LPL1-iKO, respectively), and their growth was monitored for three cycles by estimating total parasitemia at 48, 96 and 144h. **(C)** Effect of inducible knock-down of *Pf*LPL1 protein in the *Pf*LPL1-DD transgenic parasites on the development of neutral lipid body. Synchronous transgenic parasites at ring stages were grown till late trophozoite stages in presence or absence of Shld1 (control and iKO respectively) and stained with Nile red. Fluorescence images of trophozoites stage transgenic parasites in PfLPL1-iKO set, showing the reduction in Nile red fluorescence intensity and loss of GFP-fluorescence as compared to control parasite. The parasite nuclei were stained with DAPI (blue) and parasites were visualized by confocal laser scanning microscope.

**Figure S8:**
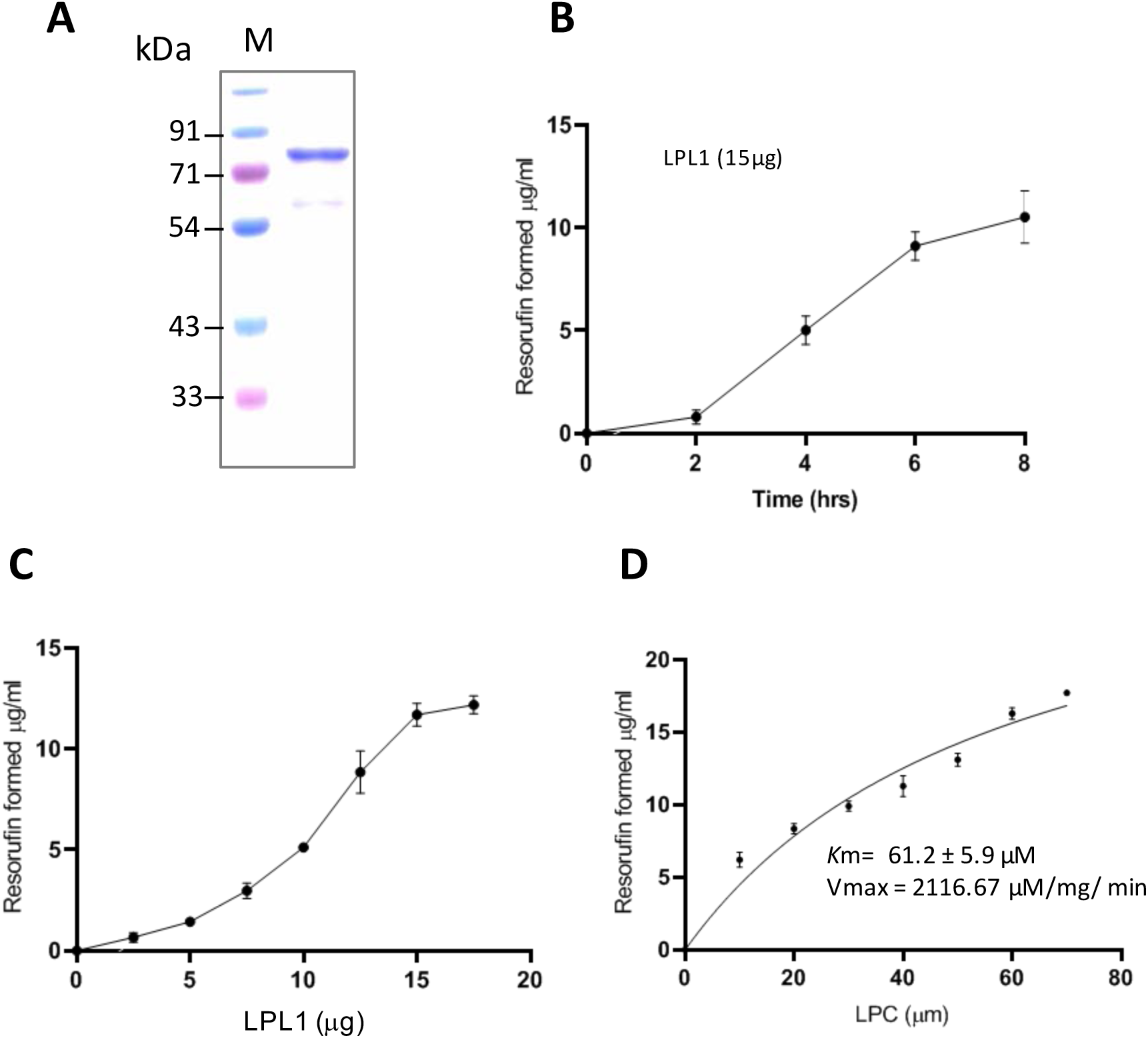
Biochemical characterization of recombinant PfLPL1: (A) Expression and purification of recombinant PfLPL1. The full gene *pflysopl1* was cloned into pETM-41 vector and recombinant protein was expressed in BL21 (DE3) *E. coli* cells. It was purified by two-step affinity chromatography using Ni^2+^-NTA and amylose resins. SDS-PAGE showing purified recombinant protein (∼72kDa). (B-D) *In vitro* activity of recombinant PfLPL1 was assessed by choline release assays using LPC as substrate and Amplex-Red detection kit. (B) Time dependent PfLPL1 activity using 15µg of recombinant protein in the assay reaction. (C) PfLPL1 activity at different protein concentrations (2.5 - 17.5µg) at 6h time point. (D) Line graph showing Michaelis-Menten fit curve developed for PfLPL1 activity using different concentrations of LPC substrate.

